# Dendritic-cell diversity in equine blood revealed by single-cell transcriptomics

**DOI:** 10.1101/2025.03.27.644174

**Authors:** Ambre Baillou, Marius Botos, Simone Oberhaensli, Iva Cvitas, Sigridur Jonsdottir, Anja Ziegler, Francisco Brito, Artur Summerfield, Eliane Marti, Stephanie C. Talker

## Abstract

Unbiased classification of equine dendritic cells (DC) is necessary to address various research questions such as the role of DC subsets in immune-mediated diseases of horses. We applied single-cell RNA sequencing (scRNA-seq) on DC enriched from the blood of two horses. All main DC subsets were detected by key gene expression, including conventional DC type 1 (cDC1; *XCR1*) and type 2 (cDC2; *FCER1A*, *CD1E*) as well as plasmacytoid DC (pDC; *TCF4*). In addition, we detected a small cluster of hematopoietic progenitors, as well as transitional DC (tDC; *FCER1A*, *TCF4*) and putative DC type 3 (DC3; *FLT3*, *CD163*). Our data confirms the previously reported phenotype of equine pDC (Flt3^+^MHC-II^low^CADM1^low^CD172a^int^), cDC1 (Flt3^+^MHC-II^high^CADM1^high^CD172a^low-int^) and cDC2 (Flt3^+^MHC-II^high^CADM1^int^CD172a^high^), while also highlighting considerable CD14 expression for cDC2. Two subclusters of equine cDC2 were found to be enriched in *FCER1A* or *CX3CR1* transcripts (cDC2.1 and cDC2.2, respectively), with suggested enhanced extravasation and T-cell stimulatory capacities of the latter. Conservation of DC subsets across species (horse, pig, human, mouse) was illustrated by enrichment analyses with subset-specific gene signatures and by cross-species data integration with publicly available scRNA-seq datasets. Our atlas of equine blood DC is a valuable resource for comparative analyses, and it forms the foundation for understanding the involvement of distinct DC subsets in infections and immune-mediated pathologies.

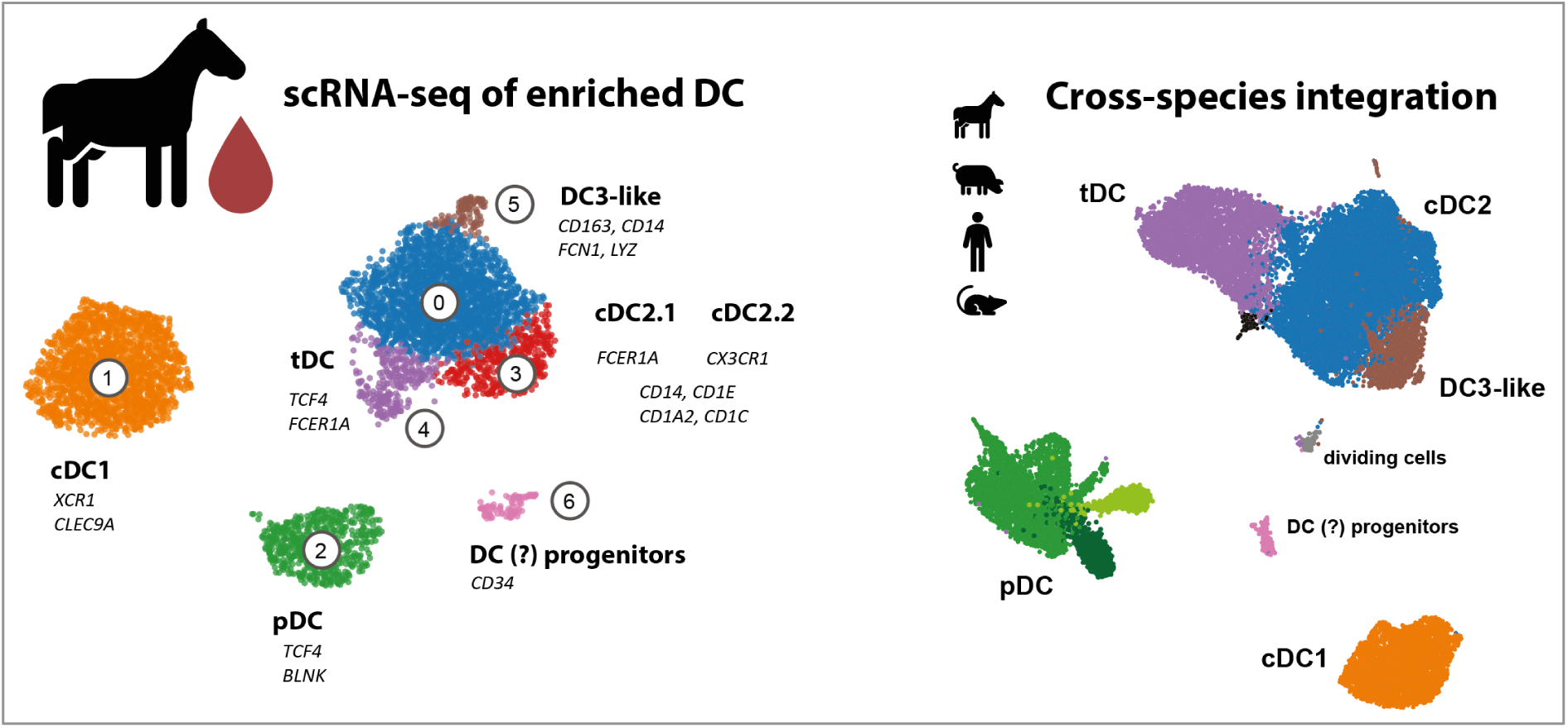

## INTRODUCTION

Dendritic cells (DC), despite their low frequency in tissues and blood, are central players in the immune system, initiating and fine-tuning adaptive immune responses (1). Traditionally, DC have been classified as conventional DC (cDC) and plasmacytoid DC (pDC). Conventional DC are grouped into two main subtypes: cDC1, specialized in cross-presenting antigens through MHC class I molecules, enabling them to prime CD8 T cells, and cDC2, known to be specialized in promoting Th2 and Th17 responses (2). While the antigen presenting capabilities of pDC are being debated, they are known for their potent antiviral effect by rapidly producing type I interferon (3), including horse (4). Flow cytometry (FCM) is widely employed for the characterization of DC subsets across species (5). However, in contrast to human and mouse, studies in veterinary species face several challenges such as limited availability of species-reactive and fluorochrome-coupled monoclonal antibodies (6). Furthermore, capturing the full spectrum of DC diversity and plasticity by a set of pre-defined markers in FCM is not feasible and prone to bias. Single-cell RNA-sequencing (scRNA-seq) has enabled the identification of novel DC subsets, including DC type 3 (DC3) (7,8) and transitional DC (tDC) in human (9), mouse (10–12), and pig (13). Across species investigated so far, DC3 have been described to share transcriptomic signatures with both cDC2 and monocytes (11,14), reflecting their origin from monocyte-dendritic-cell progenitors (7,11,15). In contrast, tDC were shown to originate from pro-pDC and to differentiate into cDC2-like cells termed tDC2, with an improved ability to prime T cells, and to produce pro-inflammatory IL-1β (12). However, delineating cDC2 subsets, monocyte-like DC, and monocytes remains a challenging task due to overlapping phenotypes and transcriptomes. In horse, a scRNA-seq study of PBMC by Patel *et al.* confirmed the identification of cDC1, cDC2 and pDC in blood (16), but, due to the small number of detected DC, could not elaborate on DC heterogeneity

Here, we describe previously unreported heterogeneity of DC in equine blood. By enriching DC prior to scRNA-seq, we were able to detect four main DC subsets (cDC1, cDC2, pDC, tDC) alongside DC3-like cells and hematopoietic progenitors / putative DC progenitors. Enrichment analysis and cross-species integration further confirmed the transcriptional similarity of DC subsets across human, pig, mouse and horse.

## RESULTS

### Equine cDC1, cDC2 and pDC defined by expression of key genes

Equine DC were enriched using FACS from freshly isolated PBMC of three horses and subsequently processed for scRNA-seq. As defined by prior phenotyping (**Supp. Fig.1A**), DC were enriched by gating on Flt3^+^ cells, and the presence of monocytes in the sorting gate was further reduced by selecting cells that expressed higher levels of CADM1 (**Figure 1A, Supp. Fig. 1B**). Upon processing, scRNA-seq data of one horse was excluded from further analyses due to low number of cells (**Supp. Fig. 2A**). As shown in **Figure 1B**, scRNA-seq of enriched DC from the remaining two horses resulted in the detection of seven clusters (clustering resolution of 0.45). By visualizing the expression of key marker genes, we identified cluster 6 as hematopoietic progenitors (*CD34*), cluster 1 as cDC1 (*XCR1*, *BATF3*, *IRF8*, *CLEC9A*, *CLNK*, *CADM1*), clusters 0, 3, 4, and 5 as cDC2-like cells (*FCER1A*, *CD1C*, *CD1A2*, *CSF1R*, *SIRPA*) and cluster 2 as pDC (*TCF4*, *BLNK*, *IRF7*, *SPIB*) (**Fig. 1B-D**). With 50% (Horse #1) and 60% (Horse #2), cDC2-like cells were found to be the most abundant cell type within enriched DC, followed by cDC1 (27% / 35%), pDC (22% / 5%) and hematopoietic progenitors (2% / 2%) **(Supp. Fig. 2B)**.

**Figure 1.**
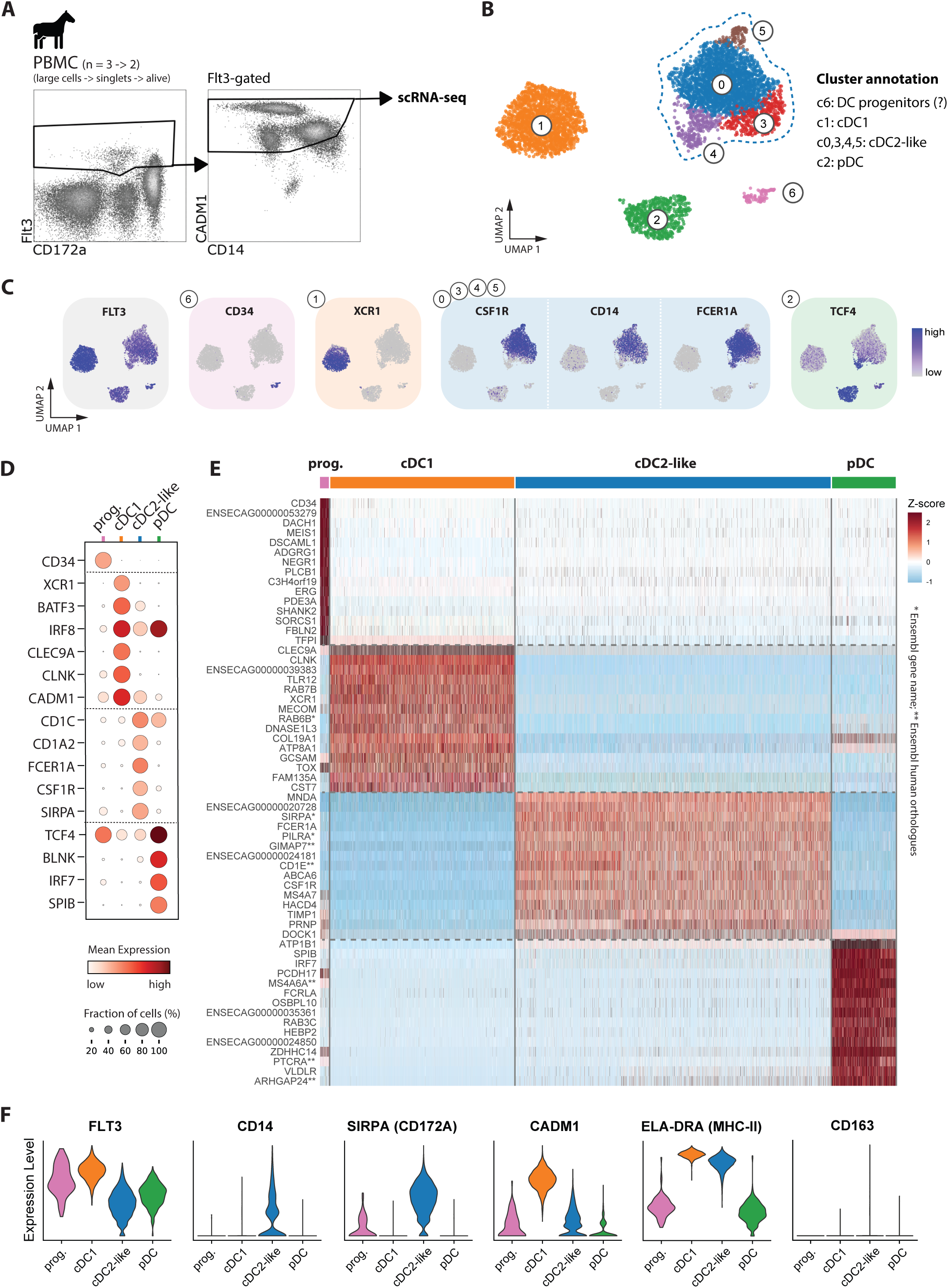
: Identification of main DC subsets in equine blood by scRNA-seq. (**A**) Gating strategy to enrich DC from equine PBMC for scRNA-seq by FACS (large cells/singlets/alive/Flt3^+^CADM1^+/hi^). (**B**) UMAP plot with annotation of putative DC progenitors, cDC1, cDC2-like cells, and pDC. (**C**) Feature plots of key markers used for subset identification. **(D)** Dot plot with additional DC subset-specific genes. (**E**) Heatmap with top 15 differentially expressed genes. Complete gene lists are given in **Supp. Table 1**. (**F**) Violin plots show transcript expression for phenotypic markers previously analyzed by flow cytometry.

Differential expression analysis (DEA) revealed additional genes with enriched transcription in hematopoietic progenitors (e.g. *MEIS1*, *DACH1, ERG, KIT, IKZF2, MSI2),* cDC1 (e.g. *CPVL*, *CST3*, *LRRK2*, *ID2, RAB7B, DNASE1L3, CD8A*), cDC2-like cells (e.g. *S100A4*, *MNDA*, *GIMAP5*, *TYROBP, IFI16, IL32*) and pDC (e.g. *CTSC*, *UGCG*, *FYB1*, *CLECL1, PLAC8B*) (**Fig. 1E, Supp. Table 1**).

Transcript detection for *FLT3, CD14, SIRPA* (*CD172a*)*, CADM1,* and *ELA-DRA* (MHC-II) in the cDC1, cDC2 and pDC clusters (**Fig. 1F**) was in line with surface protein expression detected by flow cytometry (**Supp. Fig. 1**) and with previous phenotyping reported by Ziegler *et al.* (4).

### Delineation of transitional DC (tDC) and putative DC3 from cDC2

Among cDC2-like cells, four distinct clusters were apparent (**Fig. 2A**). Two clusters (c0 and c3) were in fact comprised of cDC2, as judging from their specific expression of *CD1E, CD1A2*, and *CD1C* (**Fig. 2B**). One cluster (c4) was identified as tDC according to its expression of pDC-associated genes (e.g. *TCF4*, *SPIB*, *BLNK*) (**Fig. 2C**) alongside cDC2-associated genes (*FCER1A*, *KLF4, MNDA*) (**Supp. Table 1**). Cluster 5 displayed a monocyte-like signature (e.g. *FCN1*, *S100A12*, *C5AR1*) (**Fig. 2D**). Based on co-expression of *FLT3* (DC marker) and *CSF1R* or *CD163* (monocyte-associated) (**Fig. 2E**), we annotated this cluster as putative DC3. Gene module scoring confirmed the inflammation-associated signature of the putative DC3 cluster and additionally revealed an enrichment of complement-associated transcripts (**Fig. 2F**). Notably, cells annotated as cDC2 (c0&c3, in particular c3) were enriched in MHC-II-associated transcripts (**Fig. 2G**).

**Figure 2:**
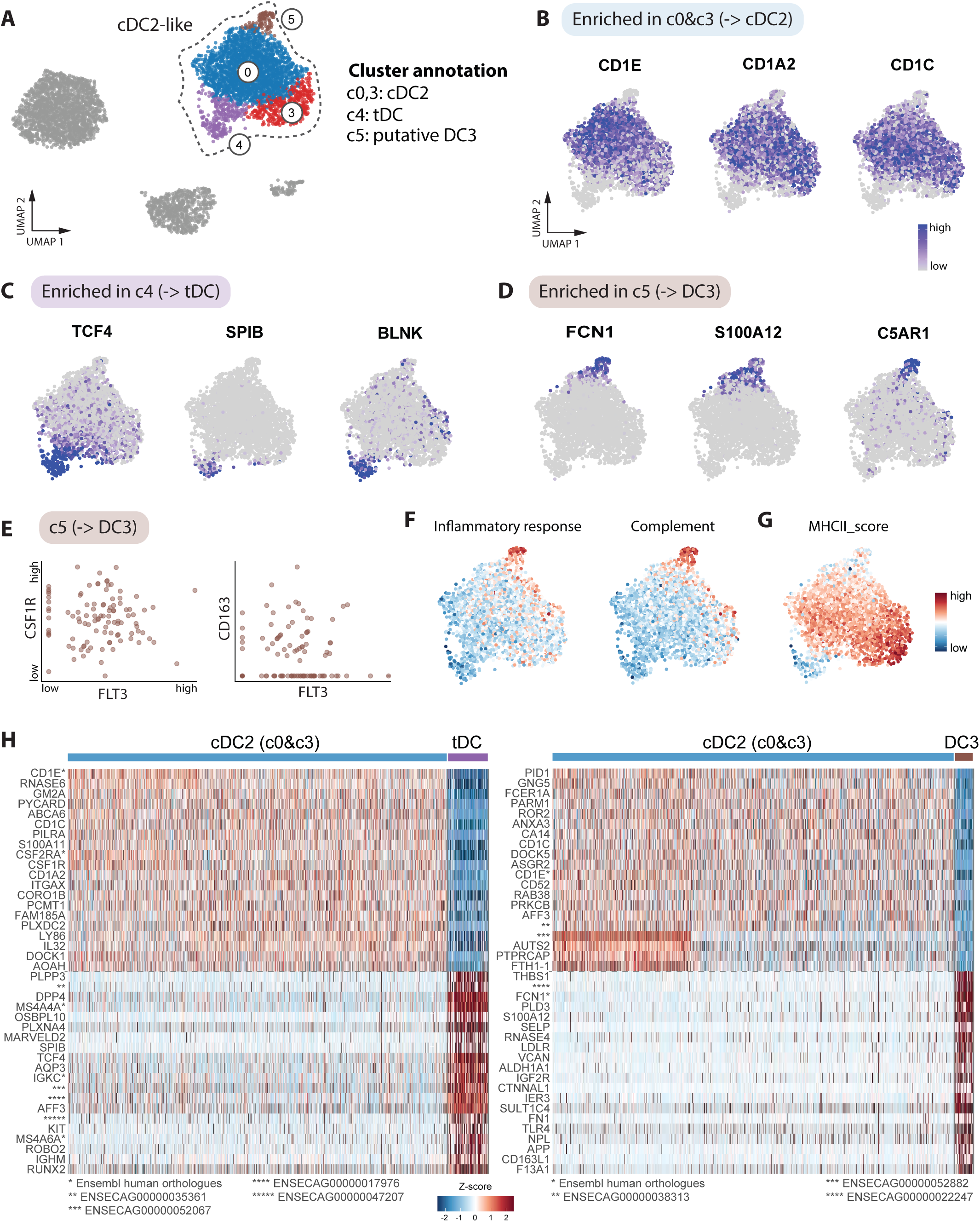
Delineation of tDC and putative DC3 from cDC2. (**A**) Annotation of cDC2-like clusters in UMAP plot. (**B**) Feature plots show expression of CD1 genes, enriched in cDC2. (**C**) Feature plots show expression of pDC-related genes, enriched in tDC. (**D**) Feature plots show expression of pro-inflammatory genes (*FCN1*, *S100A12*, *C5AR1*), enriched in DC3 (c5). **(E)** Scatter plots show co-expression of *FLT3* and monocyte markers (*CSF1R*, *CD163*) at the single-cell level in cluster 5 (DC3). (**F**) Feature plots of cDC2-like cells showing module scores for the Hallmark gene sets “inflammatory response” and “complement”. (**G**) Feature plot of cDC2-like cells showing MHC-II score (custom gene module, see **M&M**). (**H**) Heatmaps showing top 20 DEGs from pairwise comparisons between cDC2 (c0&c3) and tDC on the left, and between cDC2 (c0&c3) and DC3 on the right. Complete gene lists are given in **Supp. Table 2**.

Among the top gene transcripts enriched in tDC over cDC2 were *PLPP3* (regulator of bioactive lipids such as sphingosine-1-phosphate), *DPP4* (regulator of chemokine activity), *PLXNA4* (co-receptor for semaphorins), *CD5* (immunostimulatory receptor for effector T cell priming), *ENSECAG00000037376* (predicted ortholog of *CRLF2),* encoding a co-receptor for TSLP recently implicated in tDC2-mediated immunosuppression (17), as well as several pDC-associated genes (e.g. *RUNX2*, *IRF8*, *IKZF1*, *PLAC8B*) (**Fig. 2H, Supp. Table 2**).

In line with their monocyte-like signature, putative DC3 contained higher levels of *TLR4*, *VCAN* and *F13A1* transcripts when compared to cDC2. Other top genes differentiating DC3 from cDC2 included e.g. *THBS1*, *PLD3* and *SELP*. Complete gene lists from DEA are provided as **Supplementary Table 2**.

### Heterogeneity of cDC2

Two clusters of cDC2 were evident in our dataset (**Fig. 3A**; c0 and c3), referred to as cDC2.1 (c0) and cDC2.2 (c3). While cDC2.1 expressed higher levels of *FCER1A*, cDC2.2 were enriched in *CX3CR1* transcripts (**Fig. 3A**). Among the top differentially expressed genes in cDC2.1 (when compared to cDC2.2) were transcripts associated with pro-inflammatory (*S100A4*, *CLEC5A*, *CTSB*) and regulatory (*CD52*, *CLEC12A*) functions, as well as pattern recognition (*CLEC1A*, *CLEC5A*, *TLR2*) (**Fig. 3B-C**, **Supp. Table 2**). Moreover, cDC2.1 appeared to have high translational activity, as suggested by high expression of ribosomal genes (**Fig. 3C**).

**Figure 3:**
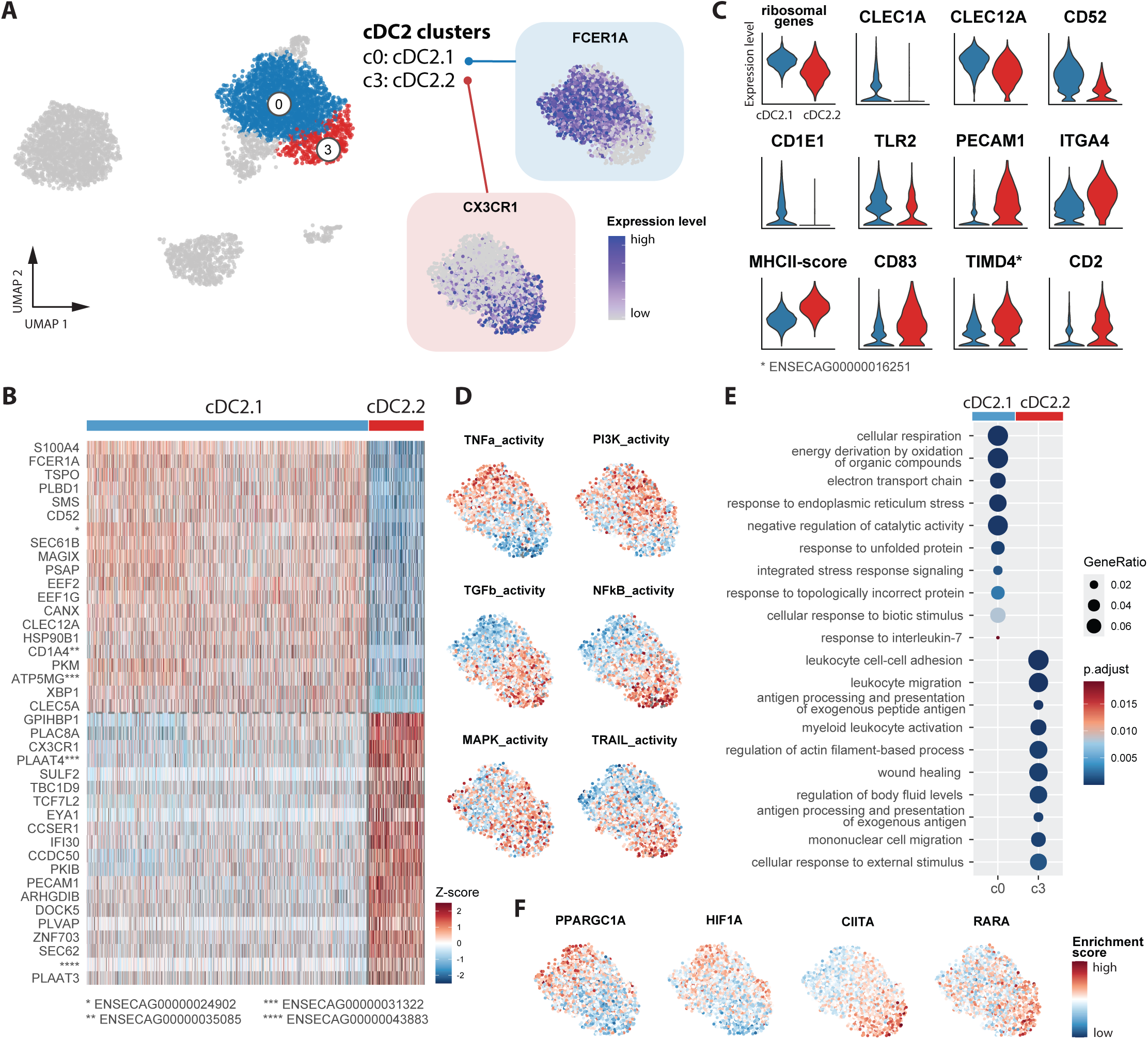
Heterogeneity of cDC2. (**A**) Annotation of cDC2 subsets (cDC1.1, cDC2.2) in UMAP plot and their expression of *FCER1A* and CX3CR1 visualized in feature plots. (**B**) Heatmap displays top 20 differentially expressed genes (DEGs) between cDC2 subsets. Complete gene lists are given in **Supp. Table 3**. (**C**) Violin plots show selected DEGs between cDC2 subsets. (**D**) Feature plots show inferred regulatory activity scores for selected PROGENy pathways across the two cDC2 clusters. (**E**) Dot plot illustrates selected enriched Gene Ontology (GO) biological process terms in the two cDC2 clusters, as determined by over-representation analysis. (**F**) Feature plots show inferred regulatory activity scores for selected differentially regulated transcription factors (TF) across the two cDC2 clusters. Complete TF list is given in **Supp. Table 3.**

In contrast, cDC2.2 stood out by enrichment of extravasation-associated transcripts such as *PECAM1* and *ITGA4* and displayed a more mature transcriptome with capacity for modulation of T-cell responses, suggested by high expression of MHC-II-associated transcripts, *CD83* and *TIMD4* (**Fig. 3B-C**, **Supp. Table 2**). Notably, cDC2.2 contained considerably higher levels of *CD2* transcripts, making CD2 a potential marker to discriminate equine cDC2 subsets by flow cytometry.

Moreover, we found differences in the activity of certain pathways (**Fig. 3D**), most notably higher TNF-ɑ activity in cDC2.1 vs. higher activity of NF-κB, MAPK, TRAIL and TGF-β in cDC2.2, with the latter two pathways potentially reflecting a more regulated state prone to apoptosis/ apoptosis induction and resolution of inflammation, altogether appearing more adapted for tissue infiltration.

Gene-ontology based over-representation analysis supported transcript enrichment for migration and antigen presentation in cDC2.2 (**Fig. 3E**). Inferred transcription factor activity further supported high metabolic activity in cDC2.1 (PPARGC1A) seemingly adapted to hypoxic environments (HIF1A), and prominent antigen presentation and T-cell stimulatory capabilities in cDC2.2 (CIITA, RARA) (**Fig. 3F**).

Notably, the signature of murine cDC2A was not clearly enriched in equine *CX3CR1*-expressing cDC2.2 compared to cDC2.1 (**Supp. Fig. 3**). This would suggest that CX3CR1 expression in equine cDC2 subsets cannot be used to differentiate putative analogs of the cDC2A and cDC2B subsets described for mouse and human.

### Delineation of DC from monocytes

Precise delineation of cDC2 and DC3 from monocytes is challenging across species. As our DC-enriched dataset likely contains very few monocytes, we took advantage of a previously published scRNA-seq dataset generated from equine PBMC (16), denoted “Patel dataset”, to integrate our dataset with a dataset containing both equine DC and monocytes (**Fig. 4A**, **Supp. Fig. 4**).

**Figure 4:**
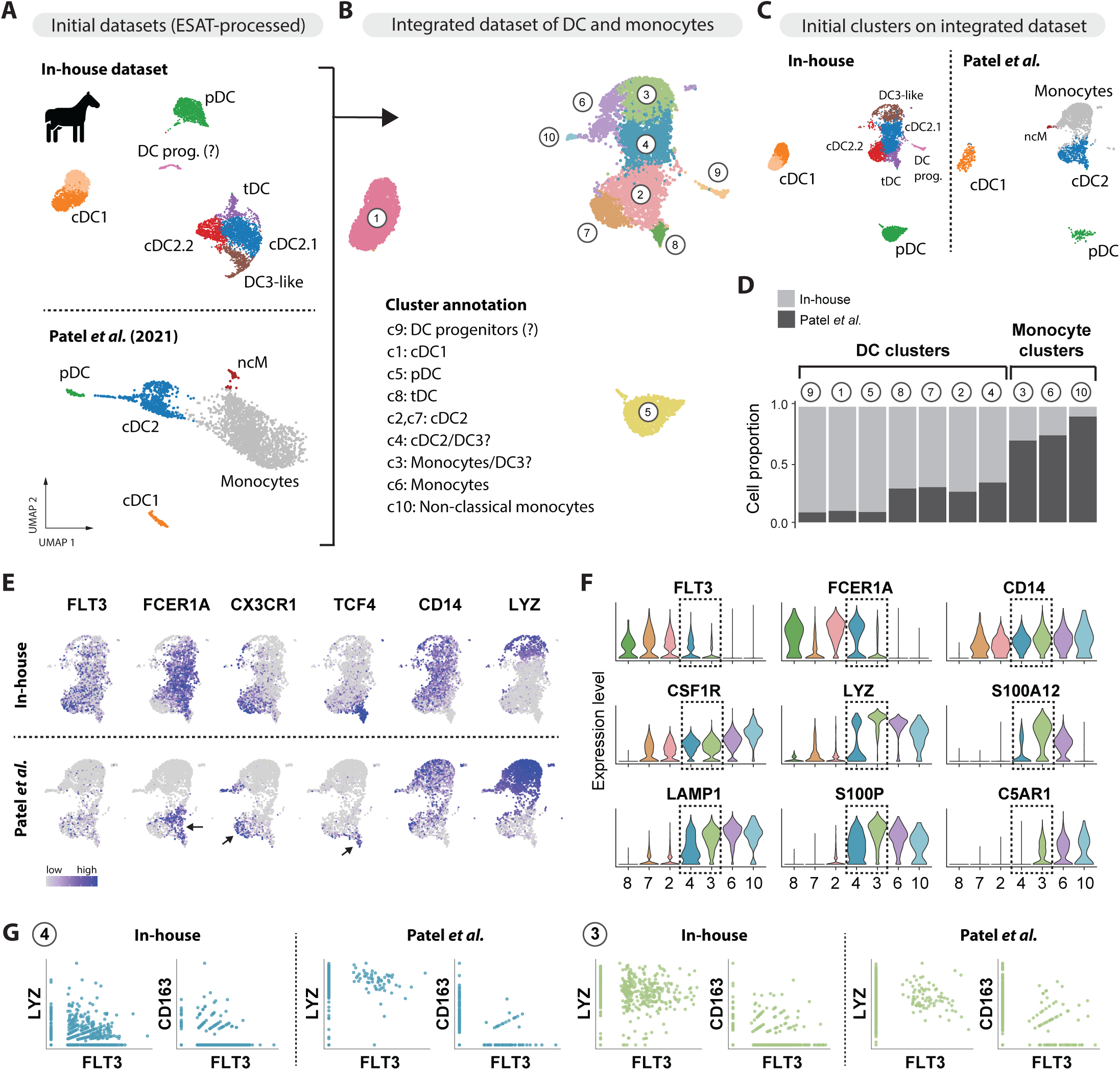
Challenges in delineating DC from monocytes. (**A**) Our in-house dataset (enriched equine DC) was re-processed according to Patel *et al.* and integrated with DC and monocytes from this previously published dataset (Patel *et al.*, equine PBMC). (**B**) UMAP plot shows annotation of ten clusters in integrated dataset. (**C**) UMAP plots show location of initial clusters (depicted in **Fig. 4A**) on integrated dataset. (**D**) Bar plots show contributions (cell proportions) of initial datasets (in-house vs. Patel *et al.*) to clusters in integrated dataset. (**E**) Key marker genes visualized in feature plots on the integrated UMAP, split by dataset (in-house vs. Patel *et al.*). (**F**) Key marker genes visualized as violin plots for clusters in integrated dataset. Clusters 4 and 3 were highlighted by dashed frames. (**G**) Scatter plots illustrate co-expression of *FLT3* and *LYZ* or *CD163* in cells of cluster 4 (blue) and cluster 3 (green) by dataset (in-house vs. Patel *et al.*).

We identified ten distinct clusters within the integrated dataset (**Fig. 4B**), with cDC1 and pDC clearly aligning across datasets (**Fig. 4C**). As expected, monocytes clustered with our DC3 (**Fig. 4C**) and originated mostly from the Patel dataset (**Fig. 4D**). As indicated by arrows in the lower panel of **Figure 4E**, cDC2 from the Patel dataset clustered according to expression of *FCER1A* and *CX3CR1* (c2 and c7), thus corresponding to the cDC2 subsets described in our study (**Fig. 3**), and according to *TCF4* expression (c8), thus corresponding to tDC described in our study (**Fig. 2**). Notably, part of the *FCER1A*-expressing cDC2 contained in our dataset (cDC2.1 in **Fig. 4C**) clustered close to monocytes (Patel dataset) and DC3-like cells (own dataset) (**Fig. 4C**), but in contrast to them almost lacked detectable *LYZ* expression (**Fig. 4E**).

Overall, while c4 and c3 appeared to be enriched for cDC2 and monocytes, respectively **(Fig. 4F)**, both of these clusters also contained DC3-like cells, as indicated by shared expression of DC-and monocyte-associated genes (**Fig. 4G**).

### Gene expression related to main DC functions

Genes associated with DC functions, such as pattern recognition, migration and adhesion, antigen presentation and T-cell modulation, as well as genes for other categories of interest, were visualized for all seven analyzed DC clusters (**Fig. 5A**).

**Figure 5:**
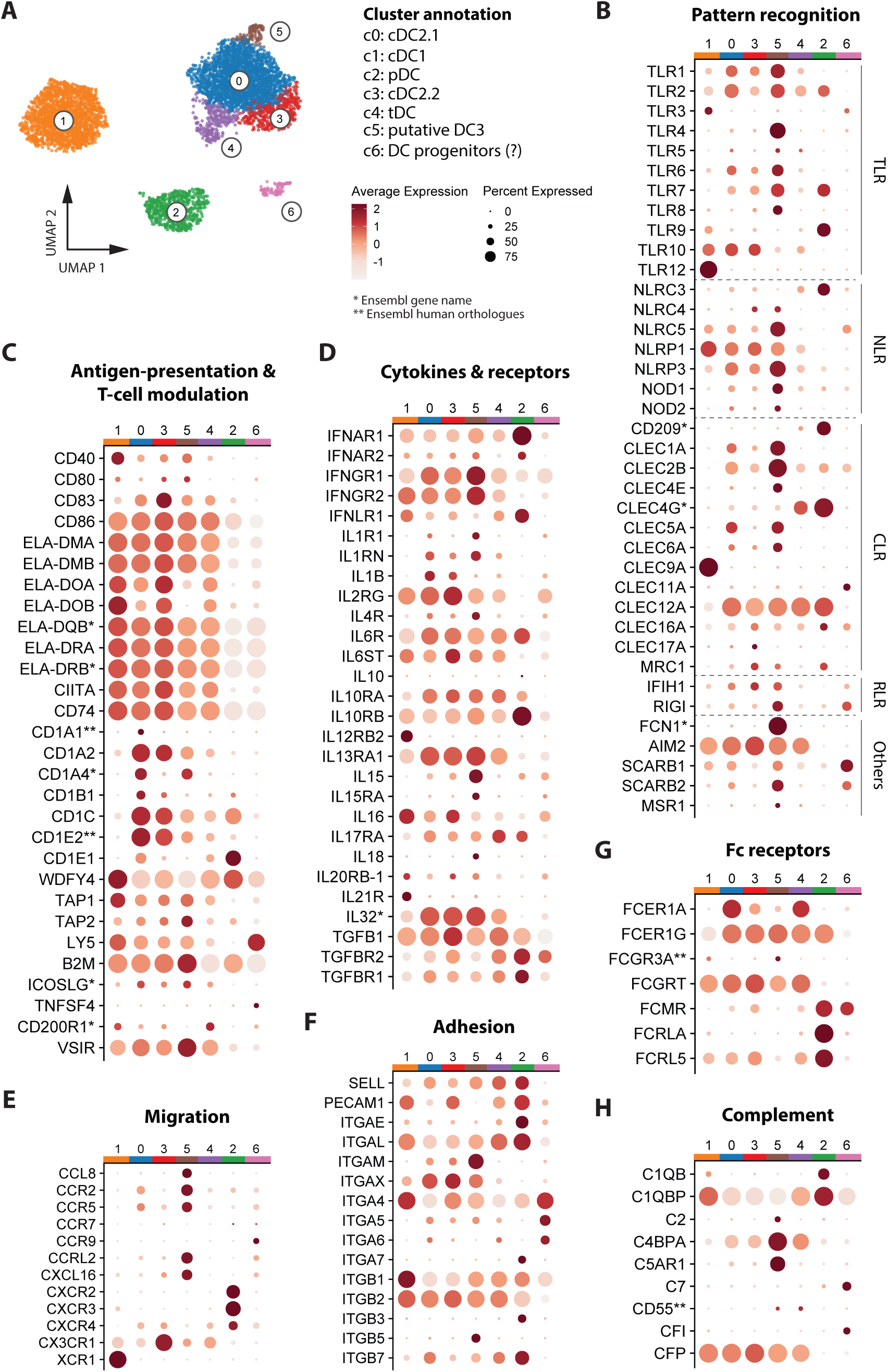
Gene expression related to main DC functions. (**A**) Cluster annotation. (**B-H**) Dot plots visualize gene expression across all annotated clusters related to pattern recognition (**B**), antigen presentation and T-cell modulation (**C**), cytokines and receptors (**D**), migration (**E**), adhesion (**F**), Fc receptors (**G**) and complement (**H**).

Among genes encoding toll-like receptors, *TLR4 and TLR8* transcripts were almost exclusively detected in putative DC3, which were also enriched for *TLR1* and *TLR6* transcripts (**Fig. 5B**). Expression of *TLR9* was selectively found in pDC, and expression of *TLR3* mostly in cDC1 and to a lesser extent in hematopoietic progenitors. Transcripts of *TLR7* were enriched in both DC3 and pDC. Most notably, cDC1 stood out by high and specific expression of *TLR12*. Overall, TLR gene expression appeared to be comparably low in tDC, with most transcripts detected for *TLR1*, *TLR2*, *TLR5* and *TLR10*. Among pattern-recognition-receptor (PRR) genes clearly expressed in tDC were *CLEC4G*, *CLEC12A* and *AIM2*, with *CLEC4G* expression exclusively shared with pDC, *CLEC12A* expression shared with all subsets except for cDC1 and progenitors, and *AIM2* expression specifically absent from pDC and progenitors.

For other PRR-associated transcripts, we found exclusive enrichment in pDC (*NLRC3*, *CD209*), strong expression in DC3 (e.g. *NLRC5*, *NLRP3*, *CLEC1A*, *CLEC2B*, *FCN1*), and exclusive or considerably high expression in progenitors (*CLEC11A*, *SCARB1*, *RIGI*).

Looking at antigen presentation and T-cell modulation, expression of MHC-II-related genes was highest in cDC1 and cDC2.2 (**Fig. 5C**), while gene expression for CD1 molecules showed a more complex pattern: cDC2 expressed the highest levels of *CD1A2*, *CD1C* and *CD1E2*, while transcripts for *CD1A4* were enriched in cDC2.1 and DC3. Notably, transcripts for *CD1E1* were enriched in pDC, which also expressed considerable levels of *CD1C*. Expression of other CD1 genes (*CD1A3*, *CD1A6*, *CD1A7*, *CD1B2*, *CD1D*) was hardly detected in the present dataset. As expected, *WDFY4* (involved in cross-presentation) showed the highest expression in cDC1. Notably, the second highest level for *WDFY4* was detected in pDC, followed by tDC and cDC2.2. Only transcripts for *CD200R1* (inhibitory receptor) were somewhat enriched (log_2_FC = 0.954) in tDC over other DC subsets.

Among genes encoding cytokines and cytokine receptors, pDC stood out by high expression of *IFNAR1*, *IFNLR1*, *IL10RB* and *TGFBR1/2* (**Fig. 5D**). Notably, cDC1 appeared to exclusively express *IL12RB2* and *IL21R* and shared high expression of *IL16* and *IL20RB-1* with cDC2.2. Expression of *IL1B* appeared to be higher in cDC2.1 than in cDC2.2, whereas cDC2.2 expressed higher levels of *IL2RG*, *IL6ST*, *IL16*, and *TGFB1* transcripts than cDC2.1. While tDC appeared to contain the highest levels of *IL17RA* transcripts, DC3 were enriched in transcripts of *IFNGR1/2*, *IL1R1*, *IL1RN*, *IL4R*, *IL15*, *IL15RA* and *IL18*.

Chemokine gene transcripts were mostly detected in DC3 (*CCL8*, *CXCL16*), alongside *CCR2*, *CCR5, and CCRL2* transcripts (**Fig. 5E**), which were also detectable in cDC2.1 at lower levels. Notably, pDC stood out by expression of *CXCR2*, *CXCR3* and *CXCR4*, while *CX3CR1* expression was highest in the cDC2.2 subset.

Looking at adhesion-associated gene expression, pDC, alongside tDC, expressed the highest levels of *SELL* (CD62L) and *ITGAL* (CD11a) (**Fig. 5F**). Notably, integrin-encoding transcripts were expressed in a highly subset-specific manner, with *ITGAE* (CD103), *ITGA7, ITGB3* and *ITGB7* clearly enriched in pDC, *ITGAM* (CD11b) and *ITGB5* in DC3, and *ITGA5* and *ITGA6* in hematopoietic progenitors.

Among genes encoding Fc receptors, tDC, alongside cDC2.1, expressed the highest levels of *FCER1A* transcripts, and shared *FCGRT* (FcRn) expression with the other cDC subsets (**Fig. 5G**). Expression of genes encoding B-cell associated Fc receptors (*FCMR*, *FCRLA*, *FCRL5*) was found to be highest in pDC. Notably, *FCMR* (encoding the Fc receptor for IgM) was the only gene in this group also clearly expressed in hematopoietic progenitors.

Looking at transcripts associated with the complement response, progenitors appeared to exclusively express *C7* (component of membrane-attack complex) and regulatory *CFI* (**Fig. 5H**), while pDC showed the highest expression of *C1QB* and *C1QBP* (initiation and regulation, respectively, of classical complement activation), and DC3 of regulatory *C4BPA* and chemotaxis/inflammation-mediating *C5AR1* (CD88). Transcripts encoding Properdin (*CFP*), binding to microbial surfaces and leading to alternative complement activation, showed the highest expression in cDC2.2 and were specifically absent from pDC and hematopoietic progenitors.

### Species comparisons support identification of equine tDC and DC3

To explore the conservation of equine DC subsets across species, we first examined the relative enrichment of DC signatures from human blood (9), murine spleen (18) and porcine blood (13) in our equine scRNA-seq clusters (**Fig. 6A**) by gene set enrichment analysis (GSEA). Results were visualized using heatmaps (**Fig. 6B-D**), complemented by feature plots to examine the distribution of enrichment scores across clusters, with manually defined thresholds for each gene set (**Supp. Fig. 5**). Overall, pDC- and cDC1-derived signatures were specifically conserved between horse and compared species, as judged by high enrichment scores for analogous subsets (**Fig. 6B-D**, **Supp. Fig. 5**).

**Figure 6:**
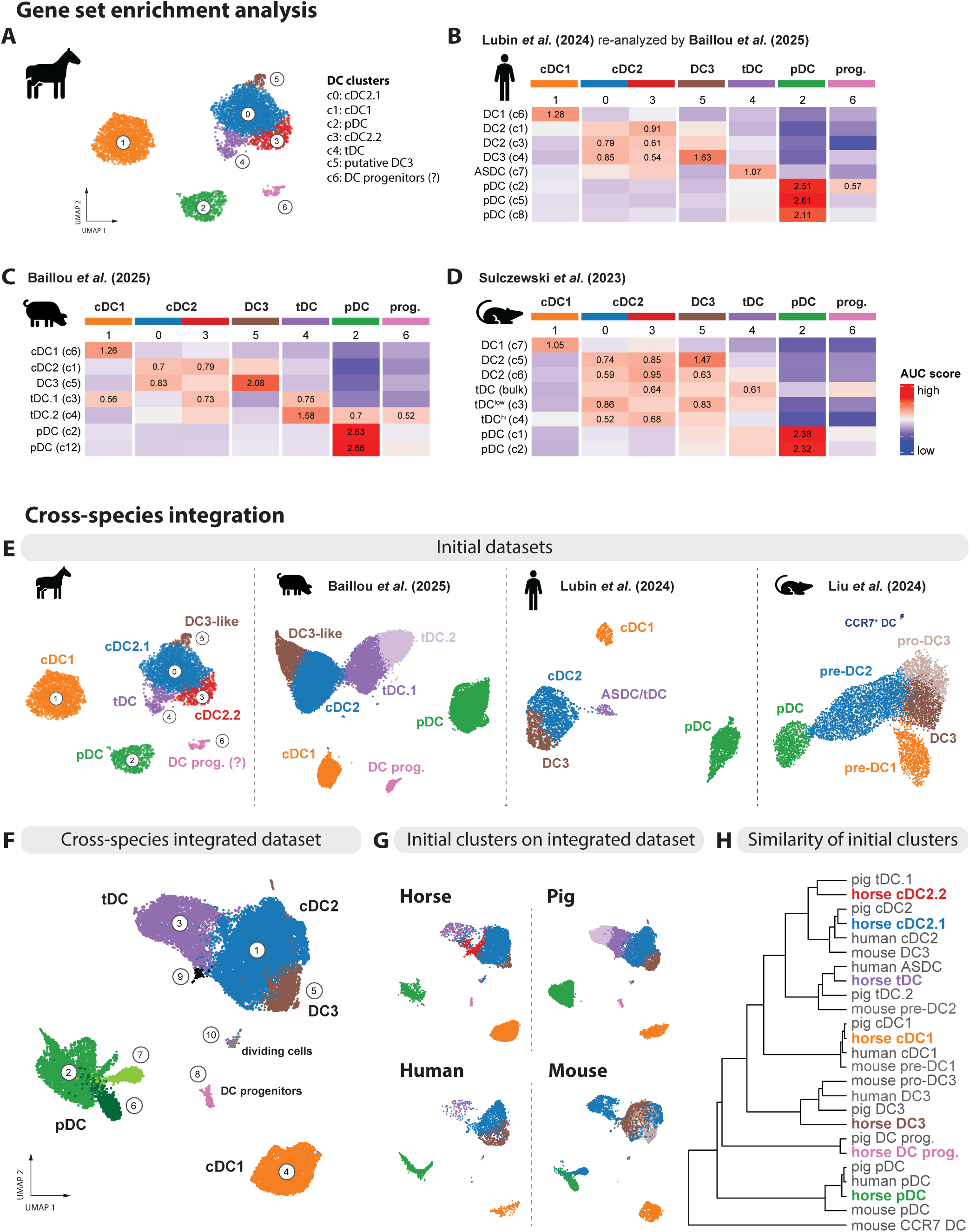
Cross-species comparison of DC subsets. (**A**) UMAP plot showing cluster annotation for equine DC subsets. (**B-D**) Heatmaps showing averaged scaled enrichment scores for gene signatures from porcine blood DC subsets (Baillou *et al.*), from murine splenic DC (Sulczewski *et al.*), and from human blood DC (Lubin *et al.*, re-analyzed by Baillou *et al.*), calculated using the AUC method (see **Supp. Table 4** for gene signatures). AUC relative scores >= 0.5 are displayed. **(D)** Integration of scRNA-seq datasets of blood-enriched DC from horse, pig (Baillou *et al.*), human (Lubin *et al.*, re-analyzed by Baillou *et al.*) and mouse (Liu *et al.*). UMAP plots show initial datasets and their respective cluster annotations. (**F**) UMAP plot showing cluster annotation for cross-species integrated dataset. (**G**) UMAP plots showing location of initial clusters (depicted in **Fig. 6E**) on integrated dataset. (**H**) Similarity tree showing correlations between the initial species-specific clusters in the integrated dataset, calculated using Euclidean distances.

Signatures from human, porcine, and murine cDC2 showed the highest enrichment in equine cDC2 clusters, with notable enrichment detected also in equine DC3, particularly for one cluster of murine cDC2.

Notably, porcine and human tDC signatures showed the highest enrichment in equine tDC. Consistent with the reported cDC2-like and tDC-like profiles of porcine tDC clusters tDC.1 and tDC.2 (13), their signatures also showed enrichment in equine cDC2.2 and pDC clusters, respectively. Moreover, the porcine tDC.1 signature was considerably enriched in equine cDC1. Murine tDC signatures were primarily enriched in equine cDC2.2 (murine tDC bulk), in equine cDC2.1 and DC3 (murine tDC^low^), or in both equine cDC2.1 and cDC2.2 (murine tDC^high^).

Porcine and human DC3 signatures resulted in the highest enrichment in the equine DC3 cluster, with additional – but comparatively lower – enrichment in the cDC2 clusters. A murine DC3 signature was not available from the mouse study we used for GSEA, but, as suggested above and further below, murine DC3 may be included within cDC2 clusters.

Next, we integrated DC-enriched scRNA-seq datasets from horse, pig, human, and mouse (**Fig. 6E**). For most species, integration resulted in co-clustering of subsets, supporting overall conservation of DC transcriptomes across species (**Fig. 6F-H**). An interesting finding was that equine cDC2.2 clustered with the porcine tDC.1 subset, echoing GSEA findings shown above, and highlighting the difficulty of clearly assigning an identity to these cell types. Notably, cells annotated as DC3 in the murine dataset clustered with cDC2 of other species (horse, pig, human), while cells annotated as pro-DC3 in the murine dataset clustered with DC3 across species. Cells annotated as pre-DC2 in the murine dataset clustered not only with cDC2 but also with tDC and pDC of the other species, likely reflecting additional heterogeneity in the previously annotated murine pre-DC2 cluster. The largest proportion of these cells annotated as murine pre-DC2 was found within the tDC cluster (c3) of the integrated dataset (**Supp. Fig. 6A**).

Expression of key genes was also largely conserved across species (**Supp. Fig. 6B**), with the notable exception of *FCER1A* and *FCN1*, which could not be detected in the murine dataset, and *F13A1*, which was highly expressed across porcine pDC and cDC2-like cells, but only detected in DC3 /proDC3 of the other species.

## DISCUSSION

Our study provides the first in-depth analysis of equine DC heterogeneity at the single-cell transcriptomic level. By performing scRNA-seq on enriched Flt3^+^ DC from blood of two horses, we identified five DC subsets, including tDC and putative DC3 alongside cDC1, cDC2, and pDC. Equine cDC1 and pDC were detected as distinct clusters expressing key genes conserved across species. Equine cDC2-like cells formed the biggest cluster (approx. 50 % within DC) and could be separated into at least two cDC2 subsets, as well as putative DC3 and tDC.

The phenotype of equine DC has previously been studied by gating on Flt3^+^ PBMC, and equine pDC were identified as Flt3^+^CADM1^low^MHC-II^low^, and confirmed by qPCR for conserved key genes (*PLAC8B*, *RUNX2*, *TCF4*) (4). Our scRNA-seq dataset validates this gating strategy and the described phenotype, with pDC expressing relatively low levels of CADM1 and MHC-II, and presumably lacking expression of CD172a and CD14. Putative cDC1 were described as Flt3^+^CD4^−^CD13^+^CD172a^−^CADM1^high^MHC-II^high^, which is in line with the high expression of *CADM1* and genes encoding MHC-II molecules in the cDC1 cluster of our scRNA-seq dataset. In the same study (4), putative cDC2 were described as Flt3^+^CD4^−^CD13^−^CD172a^high^CADM1^int^MHC-II^high^ and were gated by excluding CD14^+^ cells. As discussed below, our data (scRNA-seq and flow cytometry) clearly show CD14 expression on equine cDC2 arguing for an adjusted gating strategy without excluding CD14^+^ cells, but instead taking the expression level of CADM1 into account.

Other groups have characterized equine immune cells by scRNA-seq of PBMC (16), BALF (bronchoalveolar lavage fluid) (19–21), synovial fluid (22), and endometrium (23), but detected only low frequencies of DC. For the present study, we aimed to enrich DC as much as possible and to exclude most monocytes by stringent gating on higher CADM1 expression within the Flt3 gate. While this enhanced enrichment of DC proved beneficial to detect rare progenitors and DC subsets like tDC and DC3, our gating strategy may also have excluded some cDC2 and DC3 from our analysis, as continuous CADM1 expression levels on CD14^+^ cells did not allow for a clear separation of dendritic and monocytic cells. This phenotypic overlap at the cDC2/DC3/monocyte interface is encountered across species, making a clear separation of these subsets challenging – even at the single-cell transcriptomic level. This was also evident in our attempt to delineate DC3 and cDC2 from monocytes by integration with previously published scRNA-seq data that include equine monocytes (16). Sequencing of enriched DC alongside total mononuclear phagocytes from the same sample should minimize confounding batch effects and provide a more accurate picture of cDC2 vs. monocyte identity. However, a clear delineation of DC3 will likely remain challenging, as shown in our recently published DC atlas for blood of pigs (13).

Expression of CX3CR1 distinguishes murine T-bet^-^ cDC2 (cDC2B) from T-bet^+^ cDC2 (cDC2A) (24). At first glance, the two cDC2 clusters identified in our dataset (termed here cDC2.1 and cDC2.2) resembled CX3CR1-defined cDC2 subsets described in human and mouse (25). However, equine signatures don’t fit to the reported dichotomy of pro-inflammatory CX3CR1^+^ cDC2 and pro-resolving CX3CR1^-^ cDC2.

Rather, equine cDC2.1 (lacking *CX3CR1* expression while expressing *CCR2*) appear to be metabolically active with a relatively pro-inflammatory signature (e.g. high *IL1B* transcription), and cDC2.2 (expressing *CX3CR1* while lacking *CCR2* expression) seem to be specialized in extravasation and antigen presentation. Differential gene expression analysis revealed additional potentially interesting transcripts, such as CD52 (higher in cDC2.1) and TIMD4 (higher in cDC2.2). While CD52 expression on DC has been proposed to be downregulated upon extravasation into tissues (26), TIMD4 has been reported to be upregulated upon activation of DC and to promote T-cell expansion and survival (27) alongside efferocytosis. Although highly speculative, this would support the idea that cDC2.2 represent a more differentiated version of cDC2.1 that down-regulate CD52 while upregulating TIMD4 among other co-stimulators (e.g. CD83), ready to enter tissues, in order to support effector T cells.

Notably, CX3CR1 expression appears to also distinguish two subsets of bovine blood cDC2, as suggested by flow cytometry (28), and future scRNA-seq data of enriched bovine DC will provide a valuable resource for cross-species analyses on CX3CR1-defined cDC2 heterogeneity. Although cDC2 heterogeneity, presumably shaped by ontogeny (29,30) as well as environmental plasticity (31), will likely remain challenging to dissect – across species and tissues (32). A subset of inflammatory cDC2 described in humans (33) was later discovered to comprise a separate DC lineage: DC3, characterized by co-expression of dendritic and monocytic markers, such as CD14 (11,15). Across species, expression of CD14 is used to identify monocytes and to differentiate them from DC (2,34,35). With the discovery of *CD14*-expressing DC3 in human and mouse (7,8,11,14), this marker has been put into question. In pig, *CD14* expression appears to be similarly limited to DC3 and monocytes (13), however, our scRNA-seq data and additional phenotyping data now clearly indicate that equine cDC2 express considerable levels of CD14 and that, instead of CD14, expression of CD163 in combination with Flt3 may be suitable for distinguishing DC3 (Flt3^+^CD163^+^) from cDC2 (Flt3^+^CD163^-^) and monocytes (Flt3^-^CD163^+/-^) in horses.

Identification of tDC was based on expression of pDC-associated genes (*TCF4*, *SPIB*, *BLNK, PLAC8B*) alongside cDC2-associated genes like *FCER1A*. This transcriptomic signature shared between pDC and cDC2 is described across species investigated so far, including human (9), mouse (12), and more recently pig (13), and supports their proposed origin from pro-pDC and differentiation towards cDC2-like cells (12).

In terms of function, tDC have been described to produce pro-inflammatory IL1-β during viral infection in the mouse model (12), and to be recruited to human skin upon experimental inoculation with UV-inactivated *E. coli* (36). High levels of *IL1B* transcripts were specifically detected in equine cDC2, and to a much lesser extent in tDC. Moreover, compared to other DC subsets, overall expression of TLR genes was found to be low in equine tDC. Nevertheless, the detected gene transcripts support their involvement in bacterial recognition (*TLR1*, *TLR2, TLR5*) and regulation of inflammation (*TLR10*). In mouse models of skin inflammation and cancer, tDC-derived tDC2 were recently reported to induce regulatory T cells when sensing thymic stromal lymphopoietin (TSLP), thereby promoting immunosuppression (17). Equine tDC expressed the predicted equine ortholog of *CRLF2* (*ENSECAG00000037376*), which encodes a subunit of the TSLP receptor. This, together with prominent expression of *CD200R1*, argues for tolerogenic functions of equine tDC.

In further support of their involvement in inflammatory processes, equine tDC showed strong enrichment of *DPP4* and *IL17RA* transcripts, similar to porcine tDC (13). The peptidase CD26 (encoded by DPP4) is reported to regulate chemokine activity and thereby migration during inflammation (37), and with *IL17RA* transcripts, equine tDC appear well-equipped to respond to IL-17-rich environments, altogether suggesting their involvement in (bacterial) host defense, autoimmunity, and homeostasis of barrier tissues.

Expression of MHC-II-coding genes was lower in tDC than in cDC1 and cDC2, but considerably higher than in pDC, which is in line with transcriptomic studies performed in all three species investigated so far (12,13,18,36), and supporting their involvement in CD4 T-cell activation. Judging from the level of *WDFY4* detection, tDC may also be capable of cross-presentation. In addition, their expression of *FCGRT* (FcRn) suggests an enhanced ability to capture and present antigen-antibody immune complexes.

Moreover, equine tDC exhibited higher *CD5* expression than pDC and cDC2, mirroring observations in porcine, human and murine tDC (13,38). Notably, while CD5 may serve as a conserved marker distinguishing tDC from cDC2 and pDC across species, its expression may also indicate enhanced capabilities for T-cell priming (39).

Like porcine tDC, equine tDC were also strongly enriched in transcripts encoding Plexin 4A (*PLXNA4*), a co-receptor for class 3 semaphorins. Notably, semaphorin 3A signaling was reported to be important for DC transmigration into the lymphatic system by inducing actin-myosin contraction (40). Although Plexin A1 was described as the involved co-receptor on DC in this mouse study, Plexin A4 may serve a similar function in equine tDC for extravasating the vascular or lymphatic system. It remains to be determined if and how tDC can enter lymph nodes to engage in T-cell activation. We have preliminary data showing that porcine tDC upregulate CCR7 upon *in-vitro* TLR stimulation, which would suggest their migration to lymphatics via tissues. Before similar experiments can be performed for equine tDC, a gating strategy will need to be defined by flow cytometry. According to our transcriptomic and phenotypic data, it may be sufficient to stain CD5, alongside Flt3, CD72a, CADM1, and CD14 on equine PBMC, although their low frequency within DC may be challenging.

In our study, tDC were found to account for 4 % and 6 % of cells within DC for “Horse 1” and “Horse 2”, respectively, which is comparable to frequencies found in blood of humans and spleen of mice (9,10,12,24,36,41), and contrasting with the high frequency (approx. 30 % detected within DC) we recently reported in porcine blood (13). Future scRNA-seq studies should address differences in frequency and gene expression of tDC across tissues and species and should discuss possible implications for tDC biology. Tailored enrichment strategies will be necessary for these comparisons to reduce gating bias in FACS, as discussed above.

Cross-species mapping of DC signatures by GSEA and cross-species integration confirmed our annotation of equine DC subsets. Notably, our analyses suggest that cross-species integration can be a powerful tool to refine cluster annotations, as indicated by the dispersed clustering of murine pre-DC2 in the integrated dataset. Overall, robust cross-species comparisons of immune cells should account as much as possible for factors affecting data comparability, most notably hygienic/health status, tissue of origin and cell-enrichment strategies, among many other factors. Regarding GSEA, we must note that gene signatures (obtained by DEA) are highly dependent on the composition and clustering resolution of the initial dataset, with multiple sub-clusters of a given subset likely leading to less powerful signatures, as may be the case for signatures of murine and porcine tDC subsets used here.

Comparative analyses promise to considerably improve our understanding of newly described DC subsets. The signatures obtained from the present study are a highly valuable resource, paving the way for identifying and investigating DC subsets in lymphoid and non-lymphoid tissues of horses. Ensuing studies should explore transcriptomic perturbations of these subsets in equine immune-mediated pathologies and upon infection.

## MATERIALS AND METHODS

### Isolation of equine PBMC

Blood (100 mL) of three horses (Horse #1: male 10-year-old Freiberger; Horse #2: female 12-year-old Freiberger; Horse #3: female 8-year-old Icelandic) was collected into vacutainer collection tubes (Vacuette®; Greiner, St.Gallen, Switzerland) by puncturing the jugular vein, with sodium heparin as anti-coagulant. Peripheral blood mononuclear cells (PBMC) were isolated by density gradient centrifugation over Biocoll Separating Solution (density= 1.077 kg/l; Bioswisstec AG, Schaffhausen, Switzerland). Erythrocytes were sedimented for 30 min at room temperature (RT) before the upper leukocyte-rich plasma was collected and overlayed on Biocoll and then centrifuged at 1,800 x g for 20 min (RT). The interphase layer with PBMC was transferred to a new 50 mL tube and washed twice with PBS, with a first centrifugation for 10 min at 290 x g and a second centrifugation for 10 min at 200 x g (RT). Finally, PBMC were resuspended in PBS before cell sorting. The study was approved by the Animal Experimental Committee of the Canton of Bern, Switzerland (No. BE 2/17).

### Phenotyping of equine PBMC and DC enrichment by FACS

Freshly isolated equine PBMC were stained in 96-well U-bottom plates with 10 x 10^6^ cells per well (phenotyping) or in 5 mL FACS tubes with 80 x 10^6^ cells per tube (sorting experiments). Antibodies against CD172a, CADM1, CD14, and CD163 (phenotyping only) were used, alongside recombinant Flt3L. Bovine recombinant Flt3L-His, cross-reactive with equine Flt3, was produced in house as previously described (4,42). All incubations were performed for 20 min at 4°C, followed by washing with Cell Wash (BD Biosciences) and centrifugation at 500 x g for 5 min. Prior to the staining, all samples were incubated with equine IgG (Biorad; PEP001; 50 μg/mL) to block Fc receptors.

For phenotyping, cells were first incubated with anti-CD163 (LND68A; mIgG1) and anti-CADM1 (3E1; IgY), followed by incubation with matching secondary antibodies anti-mIgG1-PerCPeF710 (Invitrogen) and anti-IgY-biotin (Jackson ImmunoResearch). In the next step, ChromPure mouse-IgG (Jackson ImmunoResearch) was added to block remaining binding sites of the anti-mIgG1 secondary antibody, alongside Streptavidin-BV421 (BD Horizon) and Flt3L-His. In the last step, ZenonAF488-labeled anti-CD14 (105; mIgG1) was added alongside ZenonAF647-labeled anti-CD172a (DH59B; mIgG1), anti-His-PE (mIgG1; Miltenyi Biotec GmbH), and LIVE/DEAD^TM^ Fixable Near-IR stain (Thermo Fisher Scientific). For enrichment of DC by FACS, PBMC were first incubated with anti-CD172a (DH59B; mIgG1) and with anti-CADM1 (3E1; IgY), followed by incubation with corresponding secondary antibodies anti-IgG1-A647 (Invitrogen, A-21240) and anti-IgY-biotin (Jackson ImmunoResearch). Thereafter, cells were incubated with ChromPure mouse-IgG (Jackson ImmunoResearch), Streptavidin-BV421 (BD Horizon) and Flt3L-His, and in a final incubation step with ZenonAF488-labeled anti-CD14 (105; mIgG1), anti-His-PE (mIgG1; Miltenyi Biotec GmbH), and LIVE/DEAD^TM^ Fixable Near-IR stain (Thermo Fisher Scientific). Zenon labeling kits were derived from Thermo Fisher Scientific. Samples were acquired with a FACSCanto II flow cytometer (BD Biosciences) equipped with three lasers (405, 488, and 633 nm). Compensation was calculated by FACSDiva software using single-stained controls. Data were analyzed using FlowJo software (v10.10, BD Biosciences). Sorting (FACS) was performed on a MoFlo ASTRIOS EQ equipped with five lasers (355, 405, 488, 561, and 640 nm). A minimum of 30’000 cells was sorted. Cell counting and viability assessments were conducted on a TC20 automated cell counter (Biorad) using Trypan Blue dye 0.40 % (Biorad).

### Single-cell RNA sequencing

Library preparation and sequencing were done at the Next Generation Sequencing (NGS) Platform at the University of Bern. The three libraries were stored at minus 80°C, before being pooled and sequenced in one run. GEM generation & barcoding, reverse transcription, cDNA amplification and 3’ gene expression library generation steps were all performed according to the Chromium Single Cell 3’ Reagent Kits v3 User Guide (10x Genomics CG000183 Rev B) with all stipulated 10x Genomics reagents. Specifically, cell suspensions of between 700-1100 cells / µL were prepared for the GEM generation and barcoding step with a targeted cell recovery of 5’000 cells. After the GEM-reverse transcription incubation, a clean-up step was performed followed by 12 cycles of cDNA amplification. The resulting cDNA was assessed for quantity and quality using a Thermo Fisher Scientific Qubit 2.0 fluorometer with the Qubit dsDNA HS Assay Kit (Thermo Fisher Scientific, Q32854) and an Advanced Analytical Fragment Analyzer System using a Fragment Analyzer NGS Fragment Kit (Agilent, DNF-473), respectively. Thereafter, 3ʹ gene expression libraries were constructed using a sample index PCR step of 11 cycles. At the end of the protocol, an additional 0.7x bead-based clean-up of the libraries was done. The produced cDNA libraries were tested for quantity and quality using fluorometry and capillary electrophoresis as defined above. The cDNA libraries were pooled and sequenced with a loading concentration of 300 pM, paired end and single indexed, on an illumina NovaSeq 6000 sequencer with a shared NovaSeq 6000 S2 Reagent Kit (100 cycles; illumina 20012862). The read set-up was as follows: read 1: 28 cycles, i7 index: 8 cycles, i5: 0 cycles and read 2: 91 cycles. The quality of the sequencing runs was assessed using illumina Sequencing Analysis Viewer (illumina version 2.4.7) and all base call files were demultiplexed and converted into FASTQ files using illumina bcl2fastq2 conversion software version 2.19.0. An average of 543’231’771 reads/library were obtained.

### Analysis of equine scRNA-seq data

#### Genome reference and alignment

The raw sequencing data (FASTQ files) were processed using Cell Ranger v7.1 (10X Genomics) for read alignment to the equine reference genome, built with EquCab3.0 (INSDC Assembly GCA_002863925.1) as assembly and Ensembl EquCab3.0.111 as gene annotation. Filtered expression matrices from the three horses were generated using the “cellranger_count” pipeline with default arguments. Sample from “Horse 3” was excluded from further analysis (estimated number of cells: 179). Expression matrices were further analyzed in Python v3.9 (43) using scanpy v1.10.3 (44) and in R v4.3.3 (45) using mainly Seurat v5.3.0 (46), and other packages (complete list with references available on the GitHub page, see **Code availability** section).

#### Quality control and filtering

Scanpy v1.10.3 (44) was used for filtering the scRNA-seq data. Following single-cell best practices (47), we applied a median absolute deviation (MAD)-based outlier detection strategy with lenient thresholds across five quality control metrics. Accordingly, cells were excluded if they met any of the following criteria: (i) log-transformed total counts, (ii) log-transformed number of genes detected or (iii) percentage of total RNA counts from the 20 most highly expressed genes deviated more than 5 MADs above (> 5) or 3 MADs below (< 3) the median; (iv) percentage of mitochondrial gene counts > 10% or deviated beyond ± 5 MADs; and (v) percentage of hemoglobin gene counts > 5% or deviated < 8 or > 3 MADs. Thus, each sample was individually filtered and then processed for removal of doublets using scrublet v0.2.3 (48).

#### Data integration, dimensionality reduction and clustering

We applied scVI (Single-cell Variational Inference) (49) to integrate and batch-correct the data, with sample origin as a covariate. Default parameters were used, except for “n_latent = 30” and “n_layers = 2”. The scVI latent space was used for non-linear dimensionality reduction using UMAP and to construct the neighborhood graph for Leiden clustering at different resolutions.

#### Differential gene expression analysis (DEA)

We used Seurat v5.3.0 (46) for DEA on normalized and log-transformed data. The differentially expressed genes (DEGs) in each cluster were identified with the *FindAllMarkers()* function and pairwise comparisons between selected clusters were performed with the *FindMarkers()* function. Only genes expressed in at least 20% of the cells in one of the clusters being compared, and with adjusted p-value < 0.05, were selected as DEGs. For pairwise comparisons, DEGs found in 100% of the cells from both compared groups (pct.1 = 1 and pct.2 = 1) were filtered out. Complete gene lists from DEA are given in **Supplementary Table 1-3.**

#### Gene module scoring

Average expression levels of gene sets (module scores) in each cell were estimated using Seurat’s *AddModuleScore()* function. The MHC-II module scores were calculated based on seven MHC-II-coding genes (*HLA-DRA*, *HLA-DOA*, *HLA-DOB*, *HLA-DMA*, *HLA-DMB*, *ELA-DRB* and *ELA-DQB*) using “ctrl = 25” and “nbin = 15” to ensure stable expression binning and control gene selection in the equine dataset. Inflammatory response (166 genes) and complement (164 genes) modules were defined from the Hallmark gene sets of the human Molecular Signature Database (MSigDB) using msigdbr v25.1.1 (50). Human genes were mapped to equine orthologs using BioMart (Ensembl; https://www.ensembl.org/biomart/martview), retaining only one-to-one orthologous pairs, and module scores were computed with “ctrl = 50” and “nbin = 24”.

#### Pathway and transcription factor activity inference

The PROGENy and CollecTRI databases (human) were used to infer pathway and transcription factor activities, respectively, in the two equine cDC2 single-cell clusters (c0 and c3) with decoupleR v2.8.0 (51). Regulatory activity scores were estimated using the *run_mlm()* function, which applied a multivariate linear model to normalized log-transformed counts. Following score scaling, differentially regulated transcription factors (TFs) were identified with the *FindAllMarkers()* function. Only TFs with |avg_log2FC| > 1 and adjusted p-value < 0.05 were considered as differentially regulated. Complete resulting TF list is given in **Supplementary Table 3.**

#### Gene ontology over-representation analysis (ORA)

Enriched gene ontology (GO) Biological Process (BP) terms for cDC2A (c0) and cDC2B (c3) were identified using the *compareCluster()* function (fun = “enrichGO”, ont = “BP”, OrgDb = org.Hs.eg.db) from the clusterProfiler v4.10.1 package (52). Equine cluster-specific gene signatures previously defined by DEA (see **Supplementary Table 1-3**) were first mapped to their human orthologs (ENTREZ ID format) using BioMart (Ensembl). To retain the most biologically informative categories, only enriched GO BP terms corresponding to levels 4-7 in the GO hierarchy (based on ancestor counts) were included.

#### Integration with public equine dataset (PBMC-derived DC and monocytes)

Our equine scRNA-seq dataset of enriched DC (5581 cells) was integrated with DC and monocytes subsetted from a published equine PBMC dataset (2981 cells) (16). First, the PBMC data was re-processed from the publicly available count matrices generated using End Sequence Analysis Toolkit (ESAT) (53) (see **Data Availability**, and **Code Availability** for detailed pipeline). According to the initial analysis performed by Patel *et al.*, monocyte and DC clusters were selected and subclusters were characterized, allowing the identification of the main DC and monocyte subsets (**Supp. Fig. 4A-B**). A small cluster of low-quality cells (low number of genes and transcripts, high percentage of mitochondrial gene expression), as well as a cluster showing a strong neutrophil signature, were excluded from further analyses. Next, our dataset was re-processed using Ensembl v95 and ESAT (53) following the procedures originally described by the authors (16), to allow direct comparison with their dataset. After re-analysis, all previously identified DC subsets (**Fig. 1B**) were recovered based on marker gene expression profiles (**Supp. Fig. 4C**). A minor group of low-quality cells was filtered out prior to downstream analyses. Finally, both datasets were integrated as follows: (i) Each of the 9 samples (7 horses from Patel *et al.* and 2 horses from in-house dataset) were independently processed for sctransform-based normalization; (ii) Linear dimensionality reduction using PCA was performed, followed by data integration with Harmony v1.2.4 (54) selecting the sample as a covariate; (iii) Harmony embeddings were used for identifying nearest neighbors, Leiden clustering (method = “igraph”, resolution = 0.6), and non-linear dimensionality reduction using UMAP for cluster visualization in the integrated dataset (**Supp. Fig. 4D**).

#### Cross-species gene set enrichment analysis (GSEA)

Gene set enrichment analyses were performed with the AUCell package v1.24.0 (55) as initially described by Herrera-Uribe *et al.* (56) and recently adapted by Baillou et *al.* (13). The enrichment of DC subset gene signatures from various species were evaluated in the equine scRNA-seq dataset by calculating area under the curve (AUC) scores, which represent the proportion of genes from each signature that are included among the top 25% expressed genes in each cell. For each gene set, thresholds were determined manually based on AUC score distributions. (**Supp. Fig. 5**). The DC subset gene signatures were selected from four sources: (i) our recently published scRNA-seq study of porcine blood DC (10x Genomics) (13), (ii) a cDC2-focused review integrating five public murine DC scRNA-seq datasets (29), (ii) a published bulk and scRNA-seq study of murine spleen DC (10x Genomics) (12), and (iii) a published scRNA-seq dataset (10x Genomics) of human blood DC (50) that we previously re-analyzed (13). Equine one-to-one orthologs of pig, human and mouse genes were identified with BioMart (Ensembl) and selected based on the highest percentage identity to the target equine gene. Genes lacking an equine ortholog were excluded. The resulting horse-converted pig, human and murine gene signatures are provided in **Supplementary Table 4**.

#### Single-cell data integration across species using Harmony

Our equine scRNA-seq dataset (5581 cells) was integrated with three public datasets, including scRNA-seq data of blood DC population from pig (33922 cells) (13), human (2979 cells) (36) and mouse (6738 cells) (11), using Seurat v5.3.0 (46). The original porcine dataset was randomly down sampled to 10,000 cells and monocyte clusters were removed from the original murine dataset. First, human orthologs of the genes comprised in the equine, porcine and murine datasets were identified with BioMart (Ensembl). All datasets were then subsetted to their shared common genes (11,402 genes) and merged. Samples were independently processed for sctransform-based normalization (1 layer/individual or 1 layer/dataset according to the information available from each dataset), including steps of data scaling and highly variable gene identification, and for linear dimensionality reduction using PCA. For further downstream analysis, the optimal number of 30 principal components was identified by the elbow plot method. Next, data integration of the four datasets was performed with Harmony v1.2.4 (54) on PCA cell embeddings and selecting the sample origin (either individual or dataset level) and the species as covariates (batch effect correction). The resulting Harmony reduction was selected for identifying nearest neighbors, Leiden clustering (method = “igraph”, resolution = 0.4), and non-linear dimensionality reduction using UMAP for data visualization.

#### Cluster correlation analysis

Cluster correlation analysis was performed using Seurat’s *BuildClusterTree()* function on the Harmony-corrected embeddings (dims = 1:30). Euclidean distances were computed between cluster centroids, followed by hierarchical clustering (method = “average”). The resulting similarity tree was visualized using the ape v5.8-1 R package (57).

### Identification and replacement of gene identifiers

Horse gene Ensembl stable identifiers (IDs) without available gene name/symbol in the equine genome annotation file were replaced in text and figures by NCBI gene (formerly Entrezgene) accession or UniProtKB Gene Name symbol if available in the corresponding databases using BioMart (Ensembl). For genes lacking direct annotation, human orthologs were identified either through Ensembl ortholog mapping or by sequence similarity using BLAST against the human genome (GRCh38.p14). Replaced Ensembl IDs are listed in **Supplementary Table 1**. The human gene names *HLA-DOA*, *HLA-DOB*, *HLA-DMA*, *HLA-DMB* and *HLA-DRA* found in the equine genome annotation were replaced by the gene names of their equine orthologs, i.e. *ELA-DOA*, *ELA-DOB*, *ELA-DMA*, *ELA-DMB* and *ELA-DRA*, respectively.

### Preparation of Figures

Figures were prepared using FlowJo^TM^ v10.10.0 (BD Life Sciences), R v4.3.3 (45), Rstudio v2025.09.0 (58), and Adobe Illustrator v30.2.1 (https://adobe.com/products/illustrator) and Inkscape v1.3.2 (https://www.inkscape.org) softwares.

Visualization of scRNA-seq data was based on feature plots, dot plots, violin plots, scatter plots, heatmaps and histograms using Seurat v5.3.0 (46), scCustomize v3.2.0 (59), ggplot2 v3.5.2 (60), ComplexHeatmap v2.18.0 (61) and AUCell v1.24.0 (55) R packages. Heatmaps were generated with scaled and centered data (Seurat *ScaleData()* function). For improved contrast in feature plots, feature-specific contrast levels were calculated based on quantiles (q10, q90) of non-zero expression.

## DATA AVAILABILITY

Raw sequencing data from scRNA-seq of equine blood-enriched DC will be available in the European Nucleotide Archive (ENA) upon publication of the final version. Processed scRNA-seq data are available from the corresponding author upon request.

Public scRNA-seq datasets used in this study included: (i) Equine PBMC (16), available in the NCBI GEO database under the accession number GSE148416; (ii) Porcine blood DC (13), available in ENA under the accession number PRJEB101131; (iii) Murine blood DC (11), downloaded from https://doi.org/10.6084/m9.figshare.22232056.v1; and (iv) Human blood DC (36), available in the NCBI GEO database under the accession number GSE276518.

## CODE AVAILABILITY

Code and packages used for scRNA-seq data analyses will be available in a GitHub public repository upon publication of the final version.

## AUTHOR CONTRIBUTIONS

Ambre Baillou (Formal analysis, Methodology, Visualization, Writing – original draft), Marius Botos (Formal analysis, Visualization, Writing – review & editing), Simone Oberhaensli (Formal analysis, Writing – review & editing), Iva Cvitas (Data curation, Writing – review & editing), Sigridur Jonsdottir (Data curation, Writing – review & editing), Anja Ziegler (Formal analysis, Writing – review & editing), Francisco Brito (Formal analysis, Writing – review & editing), Artur Summerfield (Conceptualization, Resources, Writing – review & editing), Eliane Marti (Conceptualization, Resources, Writing – review & editing), Stephanie C. Talker (Conceptualization, Methodology, Supervision, Formal Analysis, Visualization, Writing – original Draft).

## ACKNOWLEDGEMENTS

We thank Roosheel Patel and Brad Rosenberg (Department of Microbiology, Icahn School of Medicine at Mount Sinai, USA) for ESAT processing of our dataset. Moreover, we thank Stefan Müller (FCCS, University of Bern, Switzerland) for cell sorting, the team of the NGS platform (NGSP, University of Bern, Switzerland) for single-cell RNA sequencing, and the Interfaculty Bioinformatics Unit (IBU, University of Bern, Switzerland) for access to their compute cluster.

## CONFLICT OF INTEREST

The authors declare that the research was conducted in the absence of any commercial or financial relationships that could be construed as a potential conflict of interest.

**Supplementary Figure 1: Phenotyping and enrichment of equine DC.** (**A**) Phenotyping of putative DC and monocyte subsets. Within the Flt3 gate, putative DC and monocyte subsets were defined based on expression of CD14 and CADM1. Localization of subsets in overlayed dot plot showing CD172a vs. Flt3. For each subset, expression levels of CD163 and CD172a were visualized in histograms, alongside FSC-A signal intensity. (**B**) Original gates used to enrich DC for scRNA-seq.

**Supplementary Figure 2: Quality control and cluster proportions.** (**A**) Retained cells in horses #1-3 following QC filtering. For each horse, stacked bar plots show the number of cells retained in the analysis (green), the number of cells removed due to low quality (purple; low number of reads, low number of genes, high proportion of total RNA counts that come from the 20 most highly expressed genes, high percentage of mitochondrial reads, high percentage of hemoglobin reads), and the number of cells detected as doublets and removed (orange). (**B**) Cluster annotation and proportions of cell types found in Horse #1 and Horse #2.

**Supplementary Figure 3: Enrichment of murine cDC2A and cDC2B signatures in equine cDC2.** Cross-species gene set enrichment analysis of equine cDC2 (c0&c3) using gene signatures of murine cDC2A and cDC2B derived from integrated murine blood, spleen and bone-marrow datasets (Baber *et al.*). Histograms show the distributions of enrichment scores. Threshold values (vertical red lines) were manually determined for each gene signature to distinguish non-enriched from enriched cells for visualization in feature plots. Enrichment levels among enriched cells (values > threshold) were visualized using a gradient color scale ranging from low (blue) to high (yellow) in feature plots. Heatmap visualizes enrichment scores across the two equine cDC2 clusters. Averaged, scaled AUC relative score with value >= 0.5 displayed in heatmap.

**Supplementary Figure 4: Integration of in-house and published equine datasets for delineation of DC from monocytes.** (**A**) The equine PBMC single-cell dataset from Patel *et al.* was used to subset monocytes and dendritic cells (DC). UMAP plot shows annotation of the main cell types (T cells, B cells, basophils, monocytes/DC), identified by their specific marker expression visualized in feature plots. (**B**) Monocytes and DC were selected from the PBMC dataset of Patel *et al.*, and cell subsets were identified based on key marker expression visualized in feature plots. Violin plots show quality metrics (number of genes and transcripts, percentage of mitochondrial gene expression) across clusters for identifying low-quality cells. One monocyte cluster appeared to contain neutrophils, judging from its lower gene and transcript content and specific expression of the key markers *CSF3R*, *S100A8*, *S100A9*, *E-CATH3*, and was thus removed prior to integration. (**C**) Our in-house dataset (enriched equine DC) was re-processed according to Patel *et al.*, and the previously identified DC subsets (**Fig. 1B**) were recovered based on key marker expression (shown in feature plots). Violin plots show quality metrics (number of genes and transcripts, percentage of mitochondrial gene expression) across clusters for identifying low-quality cells. (**D**) Our in-house DC dataset (excluding low-quality cells) and the monocyte/DC dataset from Patel *et al.* (excluding low-quality cells and putative neutrophils) were integrated. UMAP plot displays the annotated cell subsets in the integrated dataset, based on key marker expression visualized in feature plots.

**Supplementary Figure 5: Cross-species DC signature enrichment**. Gene signatures from porcine blood DC subsets (Baillou *et al.*; **A**), murine spleen DC (Sulczewski *et al.*; **B**) and human blood DC (Lubin *et al.*, re-analyzed by Baillou *et al.*; **C**) were used for gene set enrichment analysis of the equine scRNA-seq dataset. Histograms show the distribution of enrichment scores. Threshold values (vertical red lines) were manually determined for each gene signature to distinguish non-enriched from enriched cells for visualization in feature plots. Enrichment levels among enriched cells (values > threshold) were visualized using a gradient color scale ranging from low (blue) to high (yellow). See **Code Availability** for more details. For visualization of enrichment scores in heatmaps refer to **Fig. 6B-D**.

**Supplementary Figure 6: Cross-species DC integration**. Integration of scRNA-seq datasets of blood-enriched DC from horse, pig (Baillou *et al.*), human (Lubin *et al.*, re-analyzed by Baillou *et al.*) and mouse (Liu *et al.*) (related to **Fig. 6E-H**). (**A**) UMAP plot showing subset annotation for integrated dataset and pie charts displaying the percentage of DC subsets from each species contributing to clusters 1, 3, and 5 of the integrated dataset, normalized to their respective initial dataset sizes. (**B**) Feature plots by species showing expression of key marker genes for DC subsets in integrated dataset.

