## Supplementary Figures for "Dendritic-cell diversity in equine blood revealed by single-cell transcriptomics"

[†] *present affiliation: Next Generation Sequencing Platform and Institute for Fish and Wildlife Health, Vetsuisse Faculty, University of Bern, Bern, Switzerland*
[††] *present affiliation: Institute of Animal Pathology, Vetsuisse Faculty, University of Bern, Bern , Switzerland*
[†††] *present affiliation: Institute for Experimental Pathology, Biomedical Center, University of Iceland, Keldur, Reykjavik, Iceland and Faculty of Medicine, University of Iceland, Reykjavik, Iceland*
[††††] *present affiliation: LaboRvet AG, Reiden, Switzerland*

**Corresponding author:**
Stephanie Talker
Länggassstrasse 122, 3012 Bern, Switzerland


**Supp. Fig. 1:** Phenotyping and enrichment of equine DC (FACS).

**Supp. Fig. 2:** Quality control of scRNA-seq data and cluster proportions.

**Supp. Fig. 3:** Enrichment of murine cDC2A and cDC2B gene signatures in equine cDC2 clusters.

**Supp. Fig. 4:** Integration of in-house and published (Patel *et al.*) equine datasets for delineation of DC from monocytes.

**Supp. Fig. 5:** Cross-species DC signature enrichment (GSEA).

**Supp. Fig. 6:** Cross-species DC integration (integration of scRNA-seq datasets).

#### Supplementary Figure 1

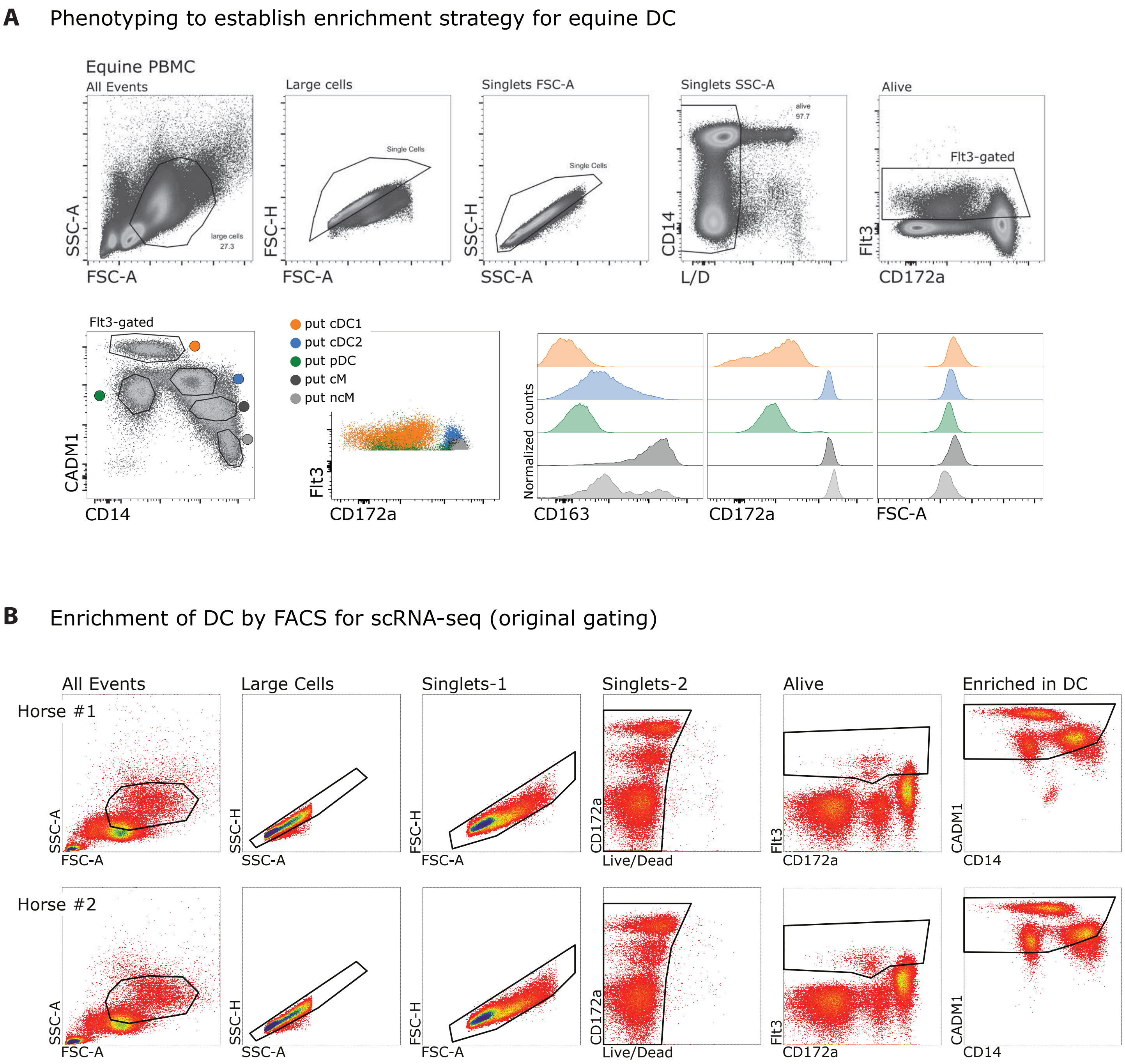


**Supplementary Figure 1: Phenotyping and enrichment of equine DC.** (**A**) Phenotyping of putative DC and monocyte subsets. Within the Flt3 gate, putative DC and monocyte subsets were defined based on expression of CD14 and CADM1. Localization of subsets in overlayed dot plot showing CD172a vs. Flt3. For each subset, expression levels of CD163 and CD172a were visualized in histograms, alongside FSC-A signal intensity. (**B**) Original gates used to enrich DC for scRNA-seq.

#### Supplementary Figure 2

**A**

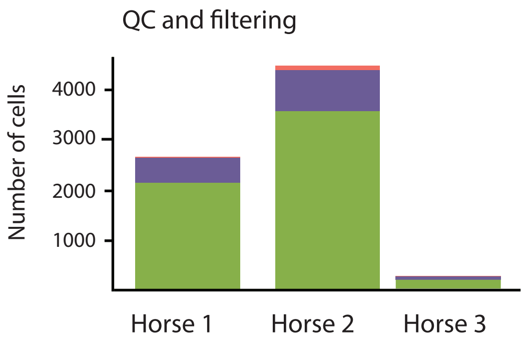


| | Breed | Age (years) | Sex | Cells after filtering scRNA-seq data |
|---|---|---|---|---|
| **Horse 1** | Freiberger | 10 | male | 2124 |
| **Horse 2** | Freiberger | 12 | female | 3537 |
| **Horse 3** | Icelandic | 8 | female | 179 (excluded from analysis) |

**B**

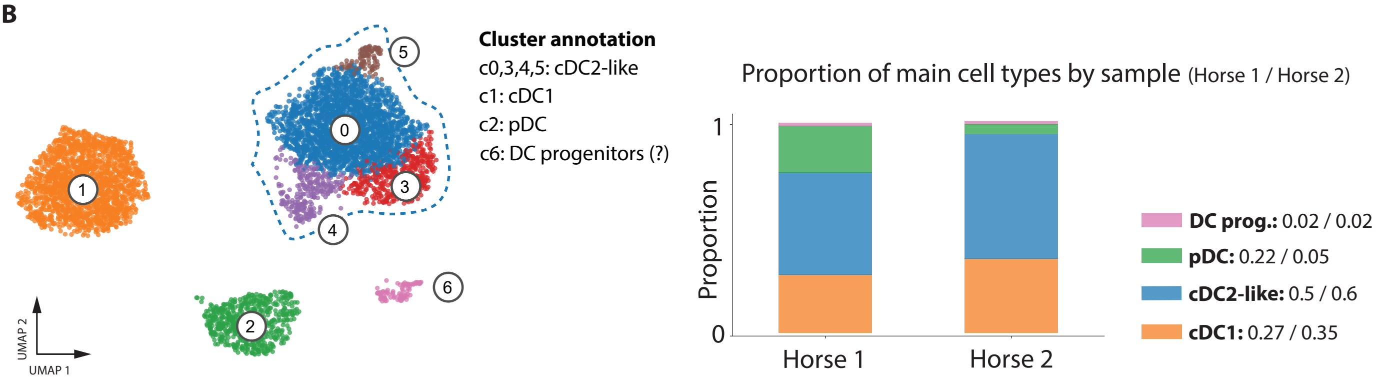


**Supplementary Figure 2: Quality control and cluster proportions.** (**A**) Retained cells in horses #1-3 following QC filtering. For each horse, stacked barplots show the number of cells retained in the analysis (green), the number of cells removed due to low quality (purple; low number of reads, low number of genes, high proportion of total RNA counts that come from the 20 most highly expressed genes, high percentage of mitochondrial reads, high percentage of hemoglobin reads), and the number of cells detected as doublets and removed (orange). (**B**) Cluster annotation and proportions of cell types found in Horse #1 and Horse #2.

#### Supplementary Figure 3

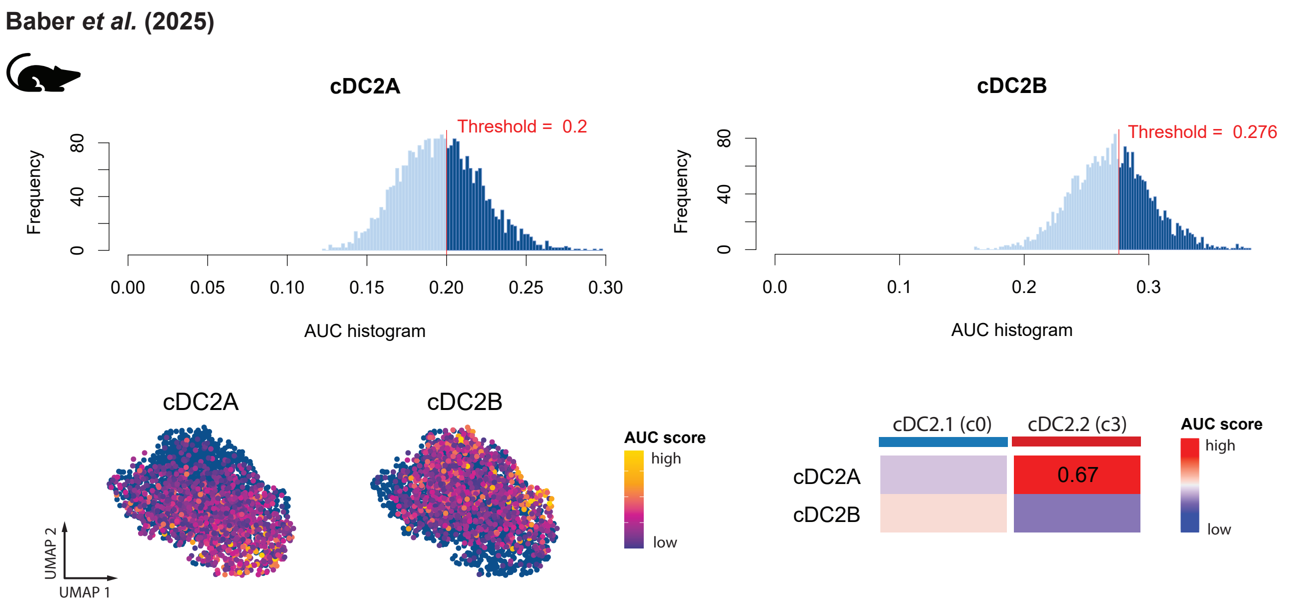


**Supplementary Figure 3: Enrichment of murine cDC2A and cDC2B signatures in equine cDC2.** Cross-species gene set enrichment analysis of equine cDC2 (c0&c3) using gene signatures of murine cDC2A and cDC2B derived from integrated murine blood, spleen and bone-marrow datasets (Baber *et al.*). Histograms show the distributions of enrichment scores. Threshold values (vertical red lines) were manually determined for each gene signature to distinguish non-enriched from enriched cells for visualization in feature plots. Enrichment levels among enriched cells (values > threshold) were visualized using a gradient color scale ranging from low (blue) to high (yellow) in feature plots. Heatmap visualizes enrichment scores across the two equine cDC2 clusters. Averaged, scaled AUC relative score with value >= 0.5 displayed in heatmap.

### Supplementary Figure 4

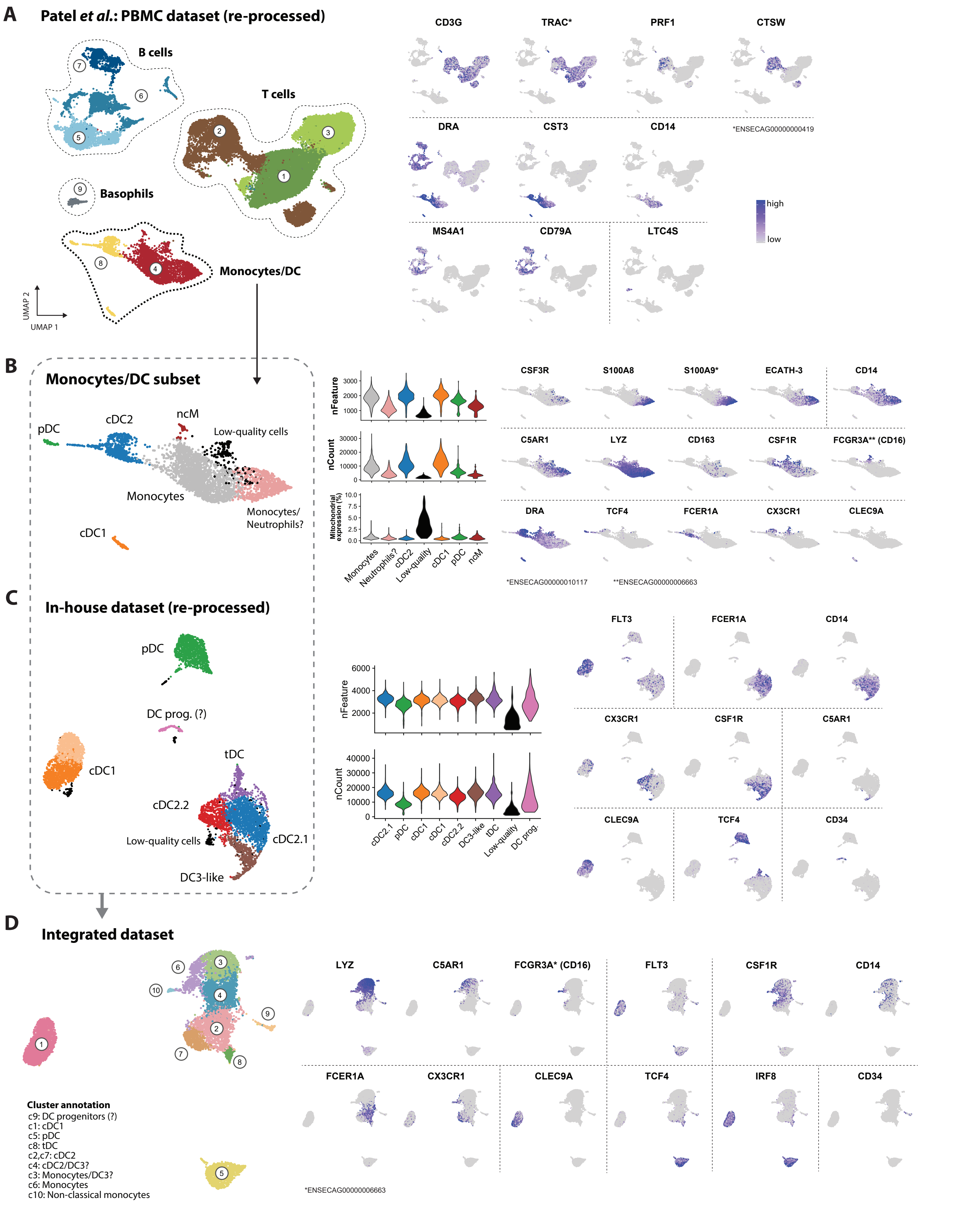


**Supplementary Figure 4: Integration of in-house and published equine datasets for delineation of DC from monocytes.** (**A**) The equine PBMC single-cell dataset from Patel *et al.* (18) was used to subset monocytes and dendritic cells (DC). UMAP plot shows annotation of the main cell types (T cells, B cells, basophils, monocytes/DC), identified by their specific marker expression visualized in feature plots. (**B**) Monocytes and DC were selected from the PBMC dataset of Patel *et al.*, and cell subsets were identified based on key marker expression visualized in feature plots. Violin plots show quality metrics (number of genes and transcripts, percentage of mitochondrial gene expression) across clusters for identifying low-quality cells. One monocyte cluster appeared to contain neutrophils, judging from its lower gene and transcript content and specific expression of the key markers *CSF3R*, *S100A8*, *S100A9*, *E-CATH3*, and was thus removed prior to integration. (**C**) Our in-house dataset (enriched equine DC) was re-processed according to Patel *et al.*, and the previously identified DC subsets (**Fig. 1B**) were recovered based on key marker expression (shown in feature plots). Violin plots show quality metrics (number of genes and transcripts, percentage of mitochondrial gene expression) across clusters for identifying low-quality cells. (**D**) Our in-house dataset (excluding low-quality cells) and the monocyte/DC-restricted dataset from Patel *et al.* (excluding low-quality cells and putative neutrophils) were integrated. UMAP plot displays the annotated cell subsets in the integrated dataset, based on key marker expression visualized in feature plots.

### Supplementary Figure 5

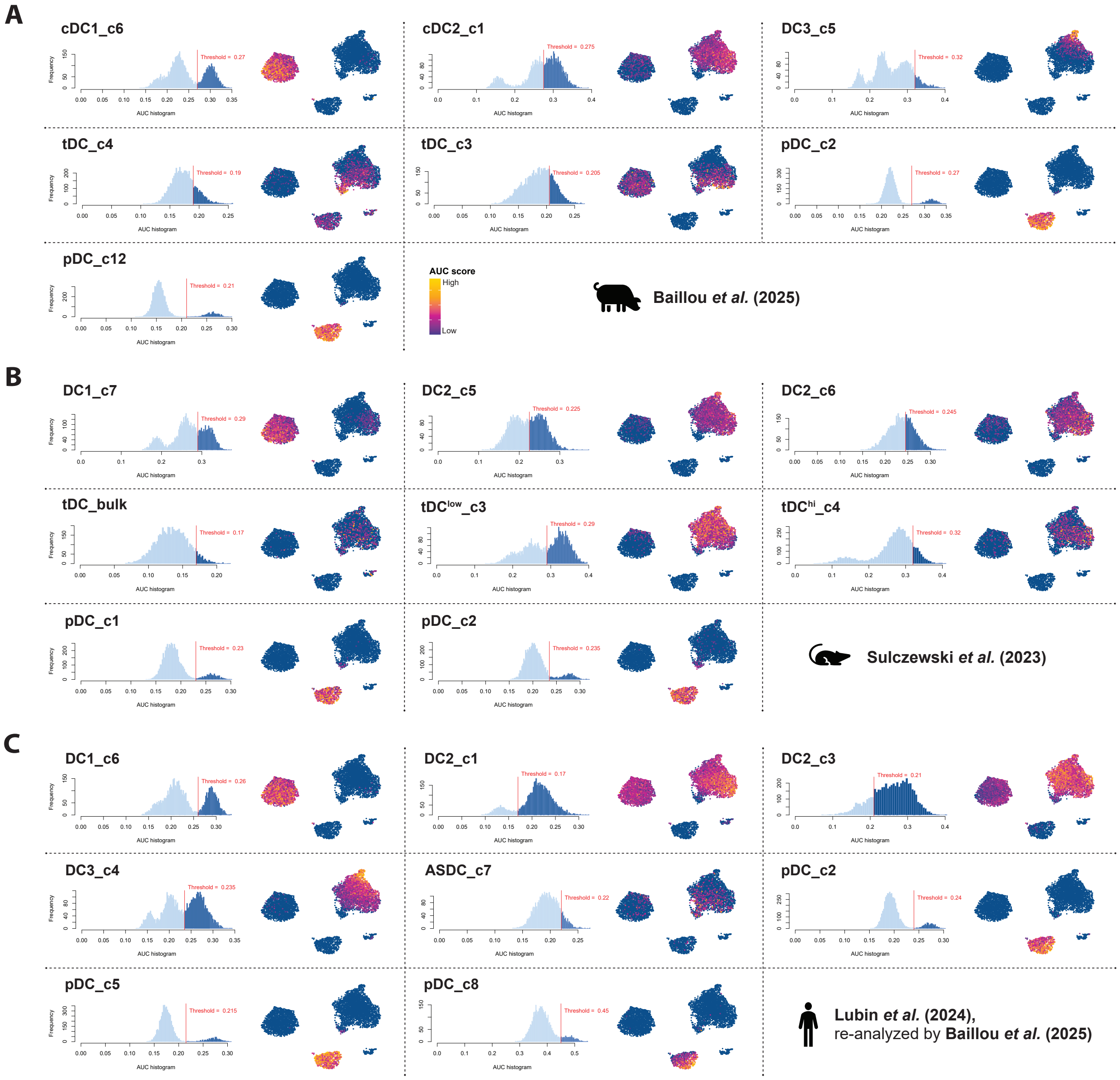


**Supplementary Figure 5: Cross-species DC signature enrichment.** Gene signatures from porcine blood DC subsets (Baillou *et al.*; **A**), murine spleen DC (Sulczewski *et al.*; **B**) and human blood DC (Lubin *et al.*, re-analyzed by Baillou *et al.*; **C**) were used for gene set enrichment analysis of the equine scRNA-seq dataset. Histograms show the distribution of enrichment scores. Threshold values (vertical red lines) were manually determined for each gene signature to distinguish non-enriched from enriched cells for visualization in feature plots. Enrichment levels among enriched cells (values > threshold) were visualized using a gradient color scale ranging from low (blue) to high (yellow). See **Code Availability** for more details. For visualization of enrichment scores in heatmaps refer to **Fig. 6B-D**.

#### Supplementary Figure 6

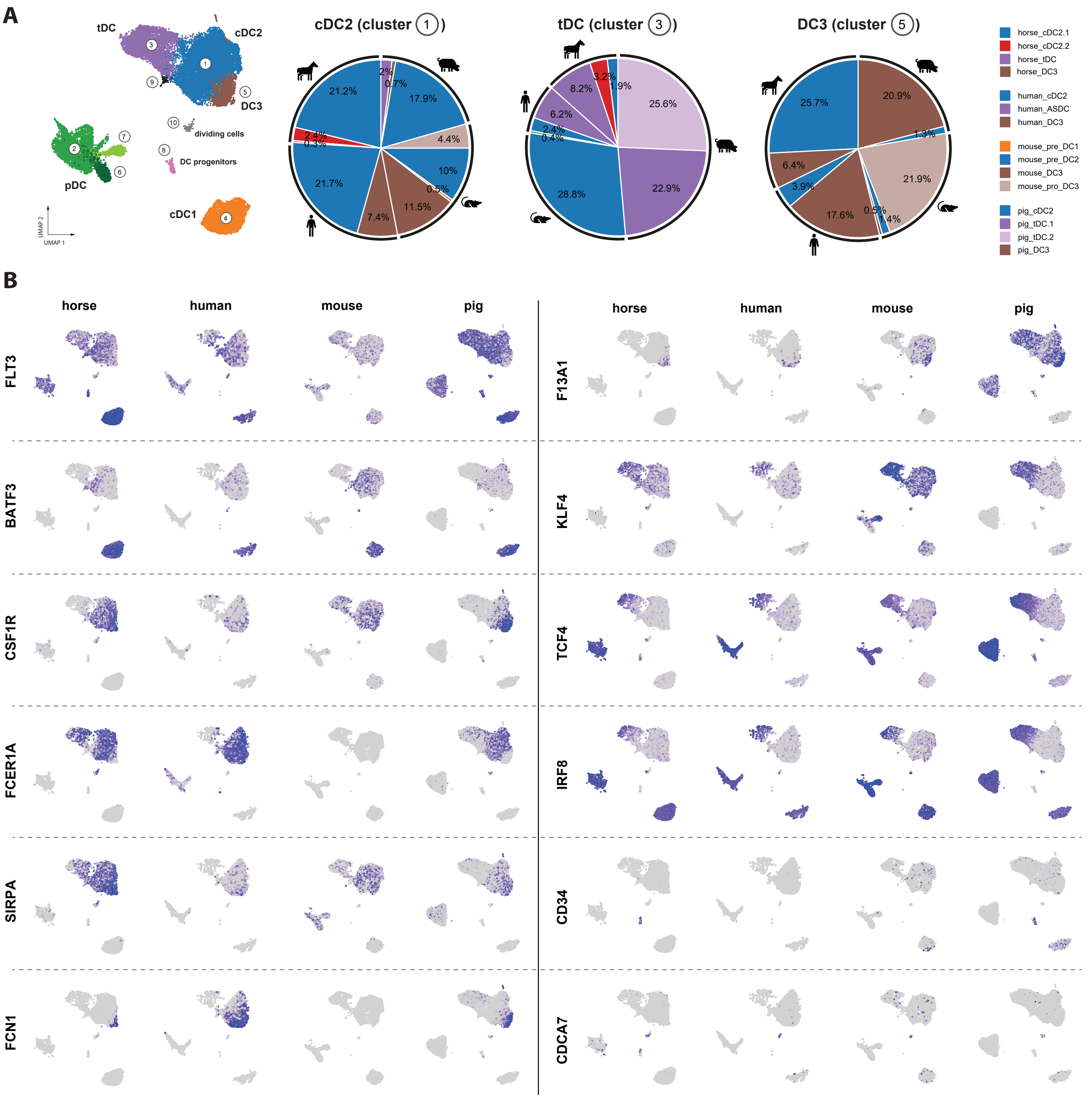


**Supplementary Figure 6: Cross-species DC integration.** Integration of scRNA-seq datasets of blood-enriched DC from horse, pig (Baillou *et al.*), human (Lubin *et al.*, re-analyzed by Baillou *et al.*) and mouse (Liu *et al.*) (related to **Figure 6E-H**). (**A**) UMAP plot showing subset annotation for integrated dataset and pie charts displaying the percentage of DC subsets from each species contributing to clusters 1, 3, and 5 of the integrated dataset, normalized to their respective initial dataset sizes. (**B**) Feature plots by species showing expression of key marker genes for DC subsets in integrated dataset.
